# Revisiting homoploid hybrid speciation in Australian cotton (*Gossypium*) species

**DOI:** 10.64898/2026.11.12.765769

**Authors:** Weixuan Ning, Mark A Arick, Joshua A Udall, Corrinne E Grover, Jonathan F Wendel

## Abstract

One of the earliest described cases of putative homoploid hybrid speciation was 35 years ago for the Australian cotton species, *Gossypium bickii*. This inference was based on discordance between chloroplast- and nuclear-based phylogenetic reconstructions, a type of incongruence that has become a relatively common observation in plants. Here we reevaluate evidence bearing on the evolutionary history of *G. bickii* in the context of an expanded sampling of all but one of the 17 described Australian cotton species, using whole-genome resequencing data based on 18,784 single-copy nuclear genes and entire plastome sequences. Genome-wide phylogenomic analyses resolved three well-defined sections (*Sturtia*, *Hibiscoidea* and *Grandicalyx*) and recapitulate the previously reported discordance between nuclear and plastid phylogenies. Although plastome inferences mirror phylogenetic discordance inferred earlier for *G. bickii* and *G. cunninghamii*, genome-wide nuclear data did not reveal compelling signals of interspecific introgression, perhaps because of population genetic nuclear swamping by a quantitatively larger paternal population and by subsequent genomic divergence. All analyses, including genome-wide phylogenetics and dating estimation, support a narrative in which there was early and rapid diversification of the three Australian sections, accompanied by early intersectional reticulation of plastomes without residual evidence in the nuclear genome.

## 1. Introduction

One of the more surprising insights of the genomics era is that interspecific gene flow is far more common in nature than recognized several decades ago (Abbott et al., 2016; Elgvin et al., 2017; Folk et al., 2018; Lamichhaney et al., 2018; Payseur & Rieseberg, 2016; Taylor & Larson, 2019). Numerous examples of homoploid reticulation leading to hybrid speciation have been proposed since the introduction of molecular markers in the last century (Abbott, 2017; Elgvin et al., 2017; Lamichhaney et al., 2018; Nieto Feliner et al., 2017; Payseur & Rieseberg, 2016; Runemark et al., 2019; Schumer et al., 2018; Taylor & Larson, 2019). These discoveries have highlighted the importance of understanding the evolutionary consequences of gene flow between divergent species, as reticulation can generate temporally stable hybrid zones, facilitate adaptive introgression, and promote colonization of novel habitats (Edelman et al., 2019; Nieto Feliner et al., 2020; Payseur & Rieseberg, 2016; Suarez-Gonzalez et al., 2018; Taylor & Larson, 2019).

Distinguishing homoploid hybrid speciation from alternative explanations for phylogenetic incongruence among molecular datasets remains challenging (Soltis & Soltis, 2009). Conceptual uncertainties surrounding the criteria for speciation (Nieto Feliner et al., 2017; Schumer et al., 2018), the potential extinction of one or more parental lineages, and the limited resolution of earlier genetic marker approaches (Alvarez & Wendel, 2003; Small et al., 2004) can all complicate the identification of hybrid speciation events. In particular, phylogenetic discordance can arise from processes such as locus-specific evolutionary histories, incomplete lineage sorting, or introgression involving unsampled ("ghost") lineages (Hibbins & Hahn, 2022; Lee-Yaw et al., 2019; Tricou et al., 2022). In addition, introgression of a chloroplast genome, for example, without survival of nuclear introgressants from the same source, may comprise a mostly neutral process that isn’t associated with adaptation, reproductive isolation, or any other criterion that might be considered “speciation”. With advances in genome sequencing technologies, reference genomes are available from throughout the tree of life, and whole genome resequencing data (WGS) generation has become more affordable (McKain et al., 2018; Ning, Meudt, et al., 2024). These advances enable genome-wide evolutionary analyses that can more reliably detect signals of hybridization and introgression (Hibbins & Hahn, 2022).

Thirty-five years ago, one of the earliest examples of homoploid hybridization in plants was described, in the pages of this journal, for the Australian cotton species *Gossypium bickii* (Wendel et al., 1991), a member of a monophyletic clade (sect. *Hibiscoidea* (G genome)) of Australian cottons (genus *Gossypium*). The 17 Australian cotton species have been phylogenetically classified into three sections, i.e., *Sturtia*, *Hibiscoidea*, and *Grandicalyx*, corresponding to the well-established “genome groups” C, G, and K, respectively (Table 1; Fig. 1; Fig. 2) (Craven et al., 1994; Fryxell, 1965; K. Wang et al., 2018), which were classically defined by comparative chromosome sizes and interspecific meiotic crossing behavior (Endrizzi et al., 1985; K. Wang et al., 2018). Among the sect. *Hibiscoidea* cottons, *G. bickii* (genome designation: G1) is one of several perennial shrubby species restricted to the arid interior of Australia (Fig. 1; Table 1). This species occurs primarily in the Northern Territory, with range extensions eastward into Queensland and westward into Western Australia. Morphologically, *G. bickii* is closely allied to the other two species of section *Hibiscoidea* (G genome), i.e., *G. australe* (G2) and *G. nelsonii* (G3), with which it shares a number of morphological synapomorphies, the most distinctive being the small facultatively cleistogamous capsules with seeds bearing short, erect epidermal trichomes (Fryxell, 1979; Stewart et al., 1987). Molecular phylogenetic reconstructions, however, indicated cytonuclear discordance in *G. bickii* (G1) implicating an evolutionary history that included inter-sectional reticulation (Seelanan et al., 1999; Wendel et al., 1991).

**Fig. 1.**
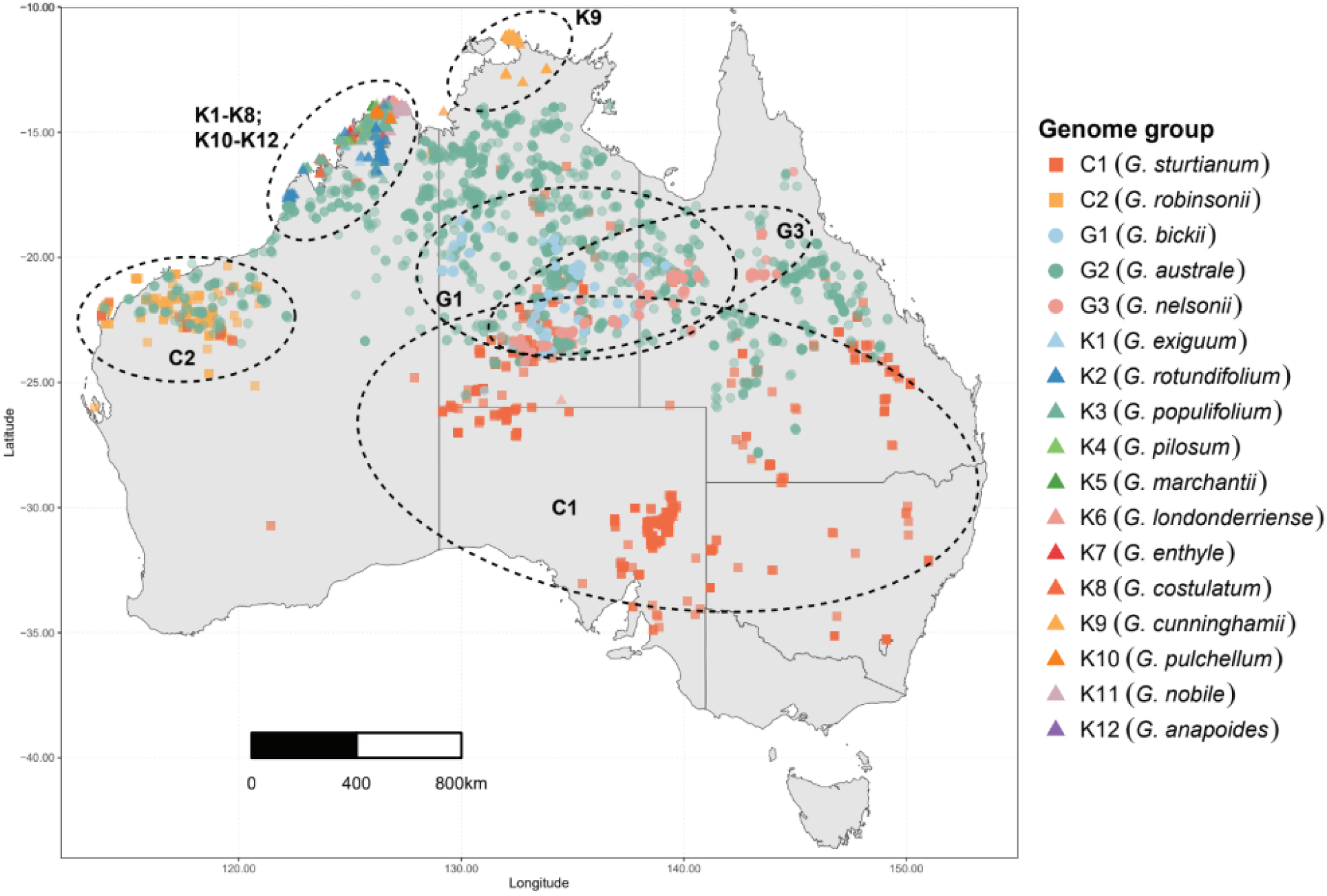
Distribution records for 17 described Australian *Gossypium* species were downloaded from GBIF (1980–2025) and plotted on Australian state boundaries from the ‘ozmaps’ R package (https://github.com/mdsumner/ozmaps). Occurrence records may contain taxonomic or georeferencing errors and were not individually validated. Species within the same genome group share the same symbol shape, whereas individual species are distinguished by color.

**Fig. 2.**
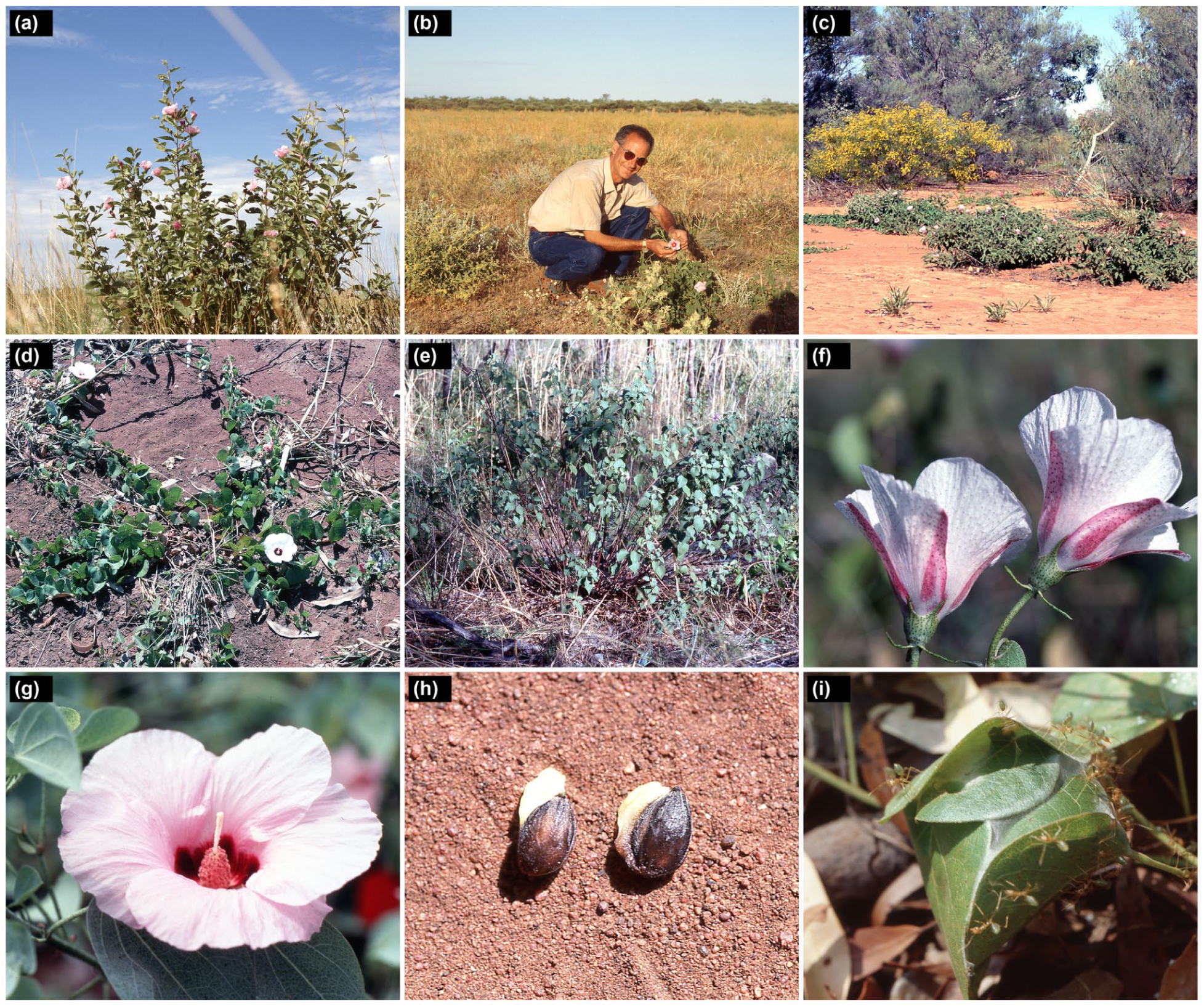
Growth forms, floral diversity, and ant-associated traits of Australian cottons (*Gossypium*). (a) *G. australe* (G2), illustrating typical shrubby habit. (b) *G. bickii* (G1; **The person pictured is J. F. Wendel, a coauthor of this manuscript.**) in flower. (c) Sprawling habit of *G. bickii* (G1) near Tennant Creek, Northern Territory, Australia. (d) Prostrate, creeping habit of *G. rotundifolium* (K2). (e) *G. pulchellum* (K10), a small, multibranched shrub. (f) Flowers of *G. enthyle* (K7). (g) Flower of *G. sturtianum* (C1), the official floral emblem of the Northern Territory, Australia. (h) Seeds of *G. rotundifolium* (K2) bearing conspicuous elaiosomes associated with ant-mediated seed dispersal. (i) Additional symbiosis with ants (*Oecophylla smaragdina*) that defend against herbivores, here illustrating nest construction on *G. marchantii* (K5).

**Table 1.**
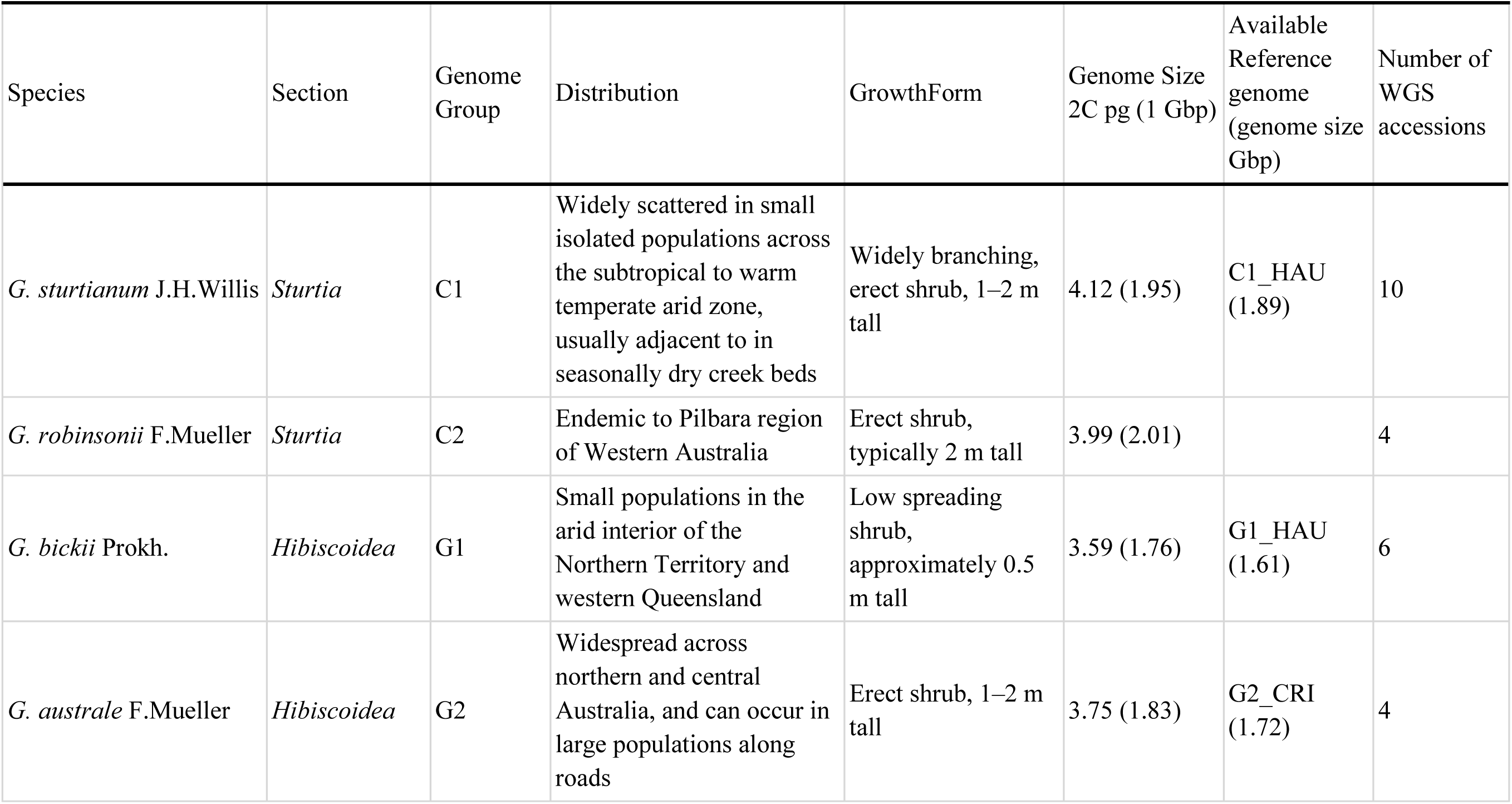
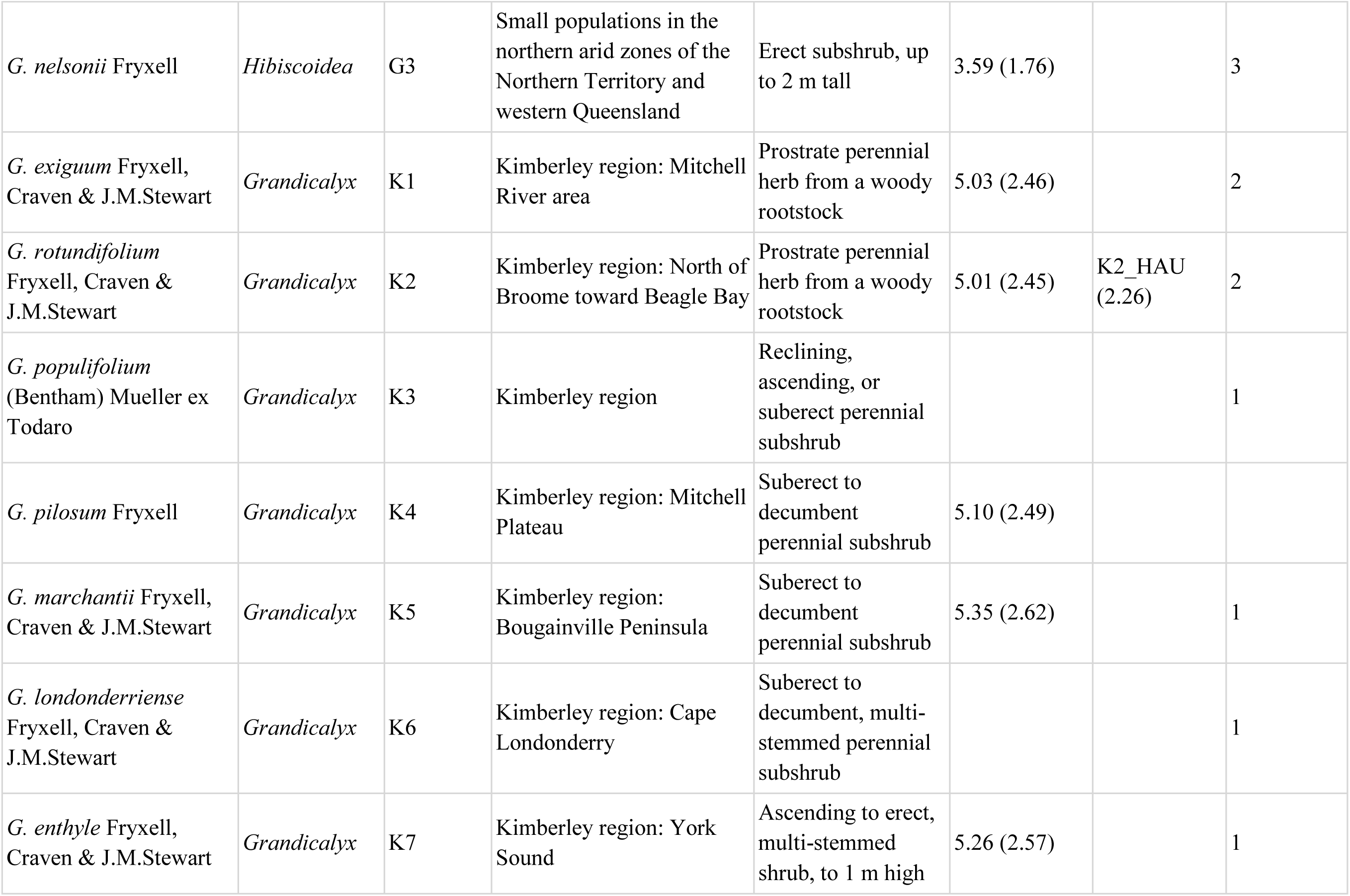
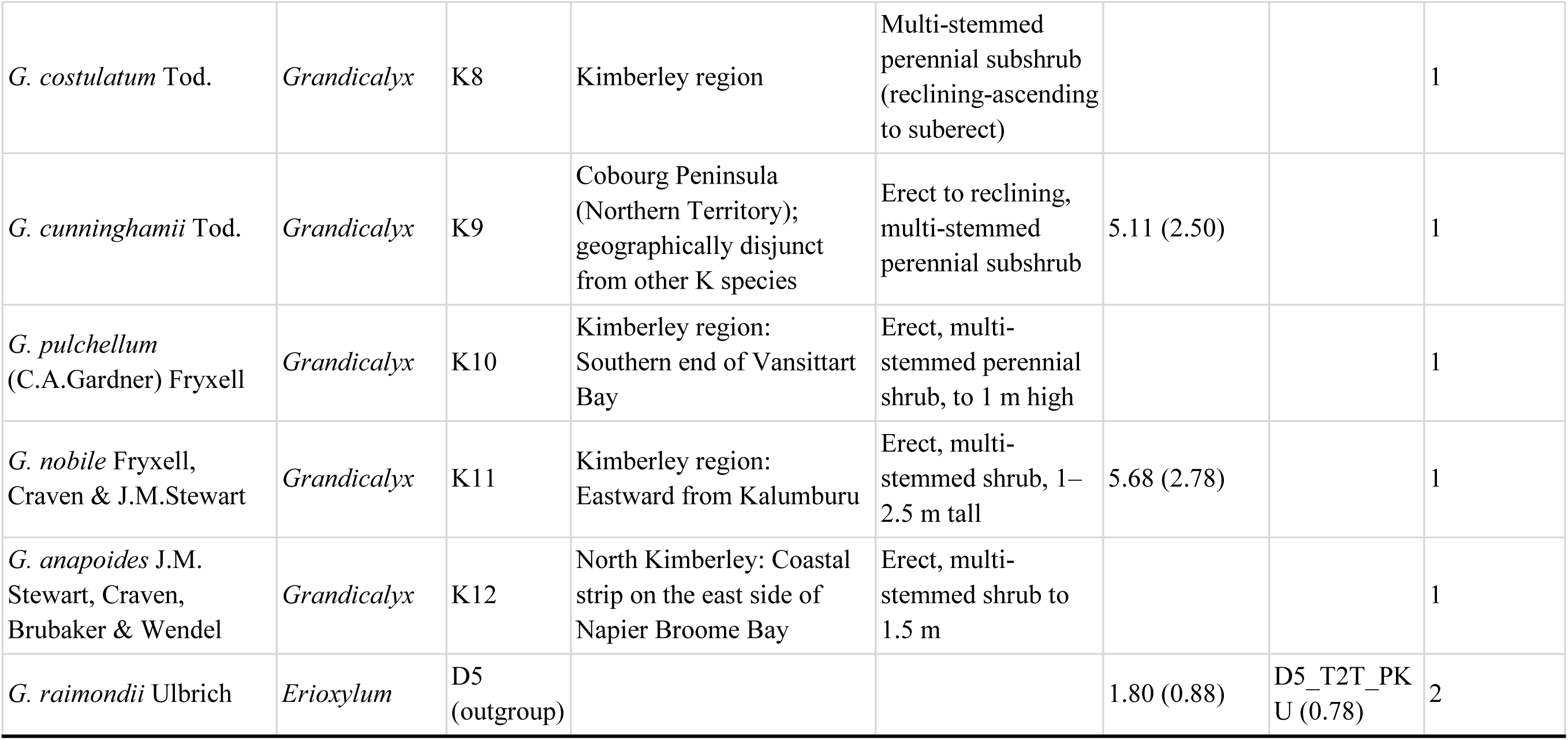
Australian cotton scientific names, sections, genome groups, distribution, growth form, available reference genomes, and number of accessions used for whole-genome resequencing. Distribution information obtained from Fryxell (1992), GBIF (https://www.gbif.org/), Australian National Botanic Gardens database (https://www.anbg.gov.au/cpbr/program/sc/cotton_mole.htm), and Craven et al., (1994); growth form information summarised from Florabase (Fryxell et al., 1992; https://florabase.dbca.wa.gov.au/; Herbarium, 1998; Stewart et al., 2015) and AusTraits (Falster et al., 2021); available reference genomes were obtained from CottonGen (J. Yu et al., 2014). Genome sizes were either classically determined (Hendrix & Stewart, 2005) and/or from genome-sequence based haploid genome sizes (1C; Mbp).

Nearly a decade later, comparative phylogenetic work based on plastid and nuclear markers revealed that a second geographically and morphologically distant species, *G. cunninghamii* (K9), also displayed evidence of historical reticulation (Seelanan et al., 1999). This latter species is from the Cobourg Peninsula of the Northern Territory, approximately 500 km NE of the remaining 11 species in section *Grandicalyx* (K genome), all of which are endemic to the widely disjunct Kimberley region (Fryxell, 1992; Seelanan et al., 1999). Species in section *Grandicalyx* (K genome) share several fire-adapted traits (Fig. 2), including a multi-stemmed herbaceous perennial growth form, a thickened woody rootstock that enables post-fire resprouting, pendent capsules that recurve after pollination and dehisce in an inverted position at maturity, and seeds bearing a lipid-rich appendage (elaiosome) that attracts ants for dispersal (myrmecochory) (Fryxell, 1992; Fryxell et al., 1992; Seelanan et al., 1999), a notable evolutionary specialization that is distinctive within the genus.

Both morphological evidence and phylogenetic analyses based on allozymes and nuclear ribosomal DNA restriction site data support the monophyly of sections *Hibiscoidea* (G genome) and *Grandicalyx* (K genome) (Craven et al., 1994; Seelanan et al., 1999; Wendel et al., 1991; Wendel & Grover, 2015); however, phylogenies based on restriction site analysis of plastid genomes revealed cytonuclear discordance relative to the nuclear phylogeny, with the geographically allopatric *G. bickii* (G1) and *G. cunninghamii* (K9) sharing a nearly identical chloroplast haplotype with *G. sturtianum* (section *Sturtia*, C1 genome). In contrast, all remaining species in sections *Hibiscoidea* and *Grandicalyx* were monophyletic with respect to chloroplast type, whereas *G. robinsonii* (C2) was unresolved in the plastid phylogeny, forming a polytomy with the other two major groups (Seelanan et al., 1999).

Phylogenetic data for the Australian *Gossypium* species remains fragmentary, being based on combinations of relatively sparse sampling of taxa and populations as well as a mix of mostly outdated cpDNA data, anonymous nuclear markers, and a small sampling of nuclear genes (Cronn & Wendel, 2004; Grover et al., 2015; Seelanan et al., 1999; Stewart et al., 2015; Wendel et al., 1991). Here we reevaluate the phylogeny of all recognized Australian *Gossypium* species using published reference genomes for each of the three taxonomic sections together with whole-genome resequencing data from 40 individuals across 16 species. Using this wealth of data, we reevaluate the phylogeny of the Australian cottons, focusing on: (1) whether whole genome resequencing data provide strong support for a particular resolution of sectional relationships; (2) genome-wide resolution of intrasectional species relationships; and (3) whether *G. bickii* (G1) or *G. cunninghamii* (K9) are supported as having evolutionary histories that include ancient introgression or homoploid hybrid speciation, as proposed in the 1990s (Seelanan et al., 1999; Wendel et al., 1991).

## 2. Materials and Methods

### 2.1 Sampling and Whole Genome Resequencing

We included all recognized species in all three taxonomic sections using material from three sources: greenhouse-maintained plants at Iowa State University, publicly available genomic resources from NCBI (https://www.ncbi.nlm.nih.gov/), and herbarium specimens from Kew (https://www.kew.org/) (Table 1; Table S1). For greenhouse-maintained plants, young leaves were collected from 18 individuals and shipped to Brigham Young University for DNA extraction using the Cetyl Trimethyl Ammonium Bromide (CTAB) method (Yuan et al., 2021), including *G. sturtianum* (6), *G. robinsonii* (2), *G. bickii* (3), *G. australe* (4), and *G. nelsonii* (3). Genomic libraries were constructed for the 18 fresh samples using the Illumina PCR-free method and sequenced on Illumina HiSeq 2500 (Brigham Young University), as reported in (Yuan et al., 2021). Additional PCR-free Illumina sequencing from 13 Australian cotton species was made available through a related project (Table S1), representing those species for which fresh tissue is unavailable. These herbarium-derived samples (collected 1967 to 1993) include *G. sturtianum* (1), *G. robinsonii* (1), and each of the 12 K genome group species, except *G. pilosum* (K4), for which no specimen was available. In addition, genome resequencing data were downloaded from NCBI for 10 Australia wild cotton individuals (Table S1), including *G. sturtianum* (3 samples; PRJNA321738, PRJNA605345, PRJNA788082), *G. robinsonii* (1; PRJNA522689), *G. bickii* (3; PRJNA321738, PRJNA605345, PRJNA788082), *G. australe* (1; PRJNA522689), *G. exiguum* (1; PRJNA321738), and *G. rotundifolium* (1; PRJNA646849). Sequencing data of two *G. raimondii* samples (D genome; sect. *Erioxylum*; Peru, South America) were also downloaded from NCBI (PRJNA202239) to be used as an outgroup for phylogenetic analysis (Wendel & Grover, 2015). In total, 43 individuals across 17 taxa, including the outgroup, were included in this study. All sequencing reads data were trimmed with Trimmomatic v.0.39 (Bolger et al., 2014) to remove adaptors sequences and low quality bases, with the following settings “ILLUMINACLIP:Adapters.fa:2:30:10:2:True LEADING:3 TRAILING:3 MINLEN:75”.

### 2.2 Selection of Single-Copy Orthologous Genes

To resolve phylogenetic relationships among Australian wild cottons, we first identified highly conserved single-copy genes shared across available reference genomes, and then applied a mapping–consensus-calling approach to extract reads corresponding to the identified orthologous gene sequences from the short-read data of each individual. Toward this end, chromosome-level genome assemblies representing four Australian wild cotton lineages were obtained from CottonGen (https://www.cottongen.org/): *G. sturtianum* (C1_HAU), *G. bickii* (G1_HAU), *G. australe* (G2_CRI), *G. rotundifolium* (K2_HAU) (M. Wang et al., 2021, 2022), and one outgroup *G. raimondii* (D5_T2T_PKU) (Huang et al., 2024). We used Liftoff v1.6.3 (Shumate & Salzberg, 2021) (“-polish -exclude_partial”) to transfer 53,167 gene annotations from the outgroup *G. raimondii* to the four Australian cotton genomes in GFF3 format and converted this to BED format using the gff2bed function in BEDOPS v2.4.40 (Neph et al., 2012). In addition, protein sequences for each reference genome were extracted from the Liftoff-generated GFF3 annotations and corresponding genome assemblies using gffread v0.12.8 (Pertea & Pertea, 2020). We then used GENESPACE v1.3.1 (Lovell et al., 2022) to identify orthologous gene sequences among the five genomes, which integrates protein homology via OthoFinder v2.5.5 (Emms & Kelly, 2019) and synteny via MCScanX (Y. Wang et al., 2012). Only conserved, single-copy orthologous gene sequences were retained. All bioinformatic pipelines and analysis settings are available on GitHub (https://github.com/Wendellab/CGKcotton).

### 2.3 Retrieving Gene Sequences from Short-Read Data

Using the gene IDs for the single-copy orthologous genes, we extracted their corresponding genomic coordinates from each of the five reference genomes (i.e., BED files with loci strand information included). These coordinates were used to retrieve reads corresponding to gene sequences from the short-read data by first mapping the short reads to the reference genomes and extracting sequences from the resulting BAM files using bam2consensus in BamBam v1.3 (Page et al., 2014), as previously done in cotton (Grover et al., 2022).

Given the diversity among Australian *Gossypium* species, we first evaluated the effect of reference genome choice on read mapping. Specifically, the trimmed reads from each individual were mapped using BWA v0.7.17 (Li, 2013) to each of the four Australian reference genomes. Mapping rates were compared across reference genomes to assess mapping performance under different reference choices. Based on mapping comparison results (see Results) and reference genome assembly qualities, we assigned *G. sturtianum* (C1_HAU) as the reference for C1 and C2 accessions, *G. bickii* (G1_HAU) for G1 and G3 accessions, *G. australe* (G2_CRI) to G2 accessions, *G. rotundifolium* (K2_HAU) for all K accessions, and the outgroup *G. raimondii* (D5_T2T_PKU) for D5 accessions. The reference-aligned BAM files were sorted (“sort”), duplicates removed (“--algo Dedup --rmdup”), and indels realigned (“--algo Realigner”) in Sentieon v202503.03 (Kendig et al., 2019).

Gene sequences for each locus were retrieved using the reference-assigned BAM files and its corresponding BED coordinates via bam2consensus with default settings. Notably, two NCBI samples with low sequencing coverage were removed, resulting in a final dataset of 40 samples. All loci were individually realigned using MAFFT v7.505 (Katoh & Standley, 2013), and then trimmed with trimAl v1.5.rev0 (Capella-Gutiérrez et al., 2009), which removes sites with a high proportion of missing data (-gt 0.7). Alignment statistics were computed for each trimmed alignment using AMAS (Borowiec, 2016), and then genes with any missing individual or non-parsimony-informative sites were excluded. We reconstructed a phylogeny for each locus using IQTree2 v2.2.2.7 (Minh et al., 2019) with 1,000 ultrafast bootstrap replicates (“-B 1000 -T AUTO”), and subsequently collapsed any internode with a bootstrap value less than 30 using nw_ed (Junier & Zdobnov, 2010) across all gene trees to improve the final species tree inference.

All gene trees were rooted using the outgroup species *G. raimondii* (D5) using reroot_trees.py (https://github.com/mossmatters/phyloscripts/tree/master/phypartspiecharts), and the final species tree was inferred using ASTRAL v.5.7.8 (Mirarab et al., 2014). We tabulated concordance between gene trees and the species tree using Phyparts (Smith et al., 2015) and plotted the results using PhypartsPieCharts (https://github.com/mossmatters/phyloscripts/tree/master/phypartspiecharts) and the R package (R. Team, 2014) ggtree (G. Yu et al., 2017).

Divergence times were estimated using Bayesian approximate-likelihood dating through the IQ2MC framework (Demotte et al., 2025). Using the rooted ASTRAL topology and the concatenated alignment of all retained loci analyzed as a single partition, IQ-TREE v3.1.3 (Wong et al., 2026) was used to estimate maximum-likelihood branch lengths under the GTR+G4 substitution model and calculate the log-likelihood gradient (first derivative) and Hessian (second derivative) required by MCMCtree for dating. Analyses used the default independent-rates, relaxed-clock settings, with ‘rgene_gamma = 2 2’, ‘sigma2_gamma = 1 10’, and ‘birth–death–sampling parameters of 1 1 0.5’. In the absence of a fossil record, a soft-bounded secondary calibration of 5 – 10 million years ago (Ma) was assigned to the common ancestor of the outgroup, *G. raimondii*, and the Australian cotton samples, as estimated previously (reviewed in Hu et al., 2021). A posterior chain was run with 1,000,000 burn-in iterations followed by 100,000 samples collected every 500 iterations.

### 2.4 Plastomes of Australian *Gossypium*

We *de novo* assembled plastomes of 36 individuals (Table S1) using GetOrganelle v1.7.7.0 (Jin et al., 2020) with default parameters (“-R 15 -k 21,45,65,85,105,125 -F embplant_pt”). For the remaining 6 herbarium-sampled individuals (all from the K genome group), the plastome did not assemble; instead, a mapping-to-consensus approach using BWA and BCFtools v1.22 (Danecek et al., 2021) was applied to extract their plastomes (Ning, Meudt, et al., 2024). Specifically, the *de novo* assembled plastome of sample K1_EL674 was selected as the reference for read mapping using BWA. BAM files containing each set of mapped reads were processed using Sentieon for sorting, duplicate removal, and indel realignment (as above). Variants were called with BCFtools, normalized (norm), filtered for gap and minimum quality (--IndelGap 5 -e ’QUAL<40’) prior to consensus reconstruction. In total, 43 plastomes were reconstructed.

Plastome phylogenies were inferred using both whole plastome sequences and cpDNA coding genes. Full cpDNA sequences were first aligned with MAFFT. Based on the GeSeq (Tillich et al., 2017) annotation of sample K1_EL674, one copy of the inverted repeat (IRB) was removed from the alignment, which then was trimmed with TrimAl (-nogaps) to exclude gapped sites. Whole chloroplast phylogenies were inferred using maximum-likelihood in IQTree2 with 1,000 bootstrap replicates (-B 1000). An additional chloroplast phylogeny was generated using only coding sequences. Chloroplast genes were extracted from the whole plastome alignment using a custom Python script (“cpdna_gene_extract.py”; https://github.com/Wendellab/CGKcotton) in conjunction with the GFF3 annotation file from K1_EL674. For genes present in duplicated regions, only one copy was retained. All gene sequences were concatenated using AMAS (concat), trimmed with TrimAl to remove gapped sites, and the resulting alignment was used for phylogenetic reconstruction in IQTree2.

### 2.5 SNP Calling from Divergent Australian Cotton Lineages

Paired-end reads of each accession were mapped to their the closest reference genome using BWA (e.g., C1_HAU to C1 and C2, G1_HAU to G1, G2_CRI to G2 and G3, K2_HAU to K genome group, and D5_T2T_PKU to D5) and genotyped in Sentieon. Each sample followed standard genotyping practices: removal of duplicated reads (“--algo Dedup --rmdup”), indel realignment (“--algo Realigner”), and base quality score recalibration (“--algo QualCal”) prior to sample genotyping (i.e. GVCF creation). From those, five VCF files (one per reference genome) were generated containing both variant and invariant sites (“--algo GVCFtyper --emit_mode all”). Indels were subsequently removed (“--remove-indels”) from all VCFs using VCFtools v.0.1.16 (Danecek et al., 2011), and only biallelic (“--min-alleles 2 --max-alleles 2”) and invariant sites (“--max-maf 0”) were remained for downstream analysis.

Because each VCF was generated using a different genome, the coordinates were transferred to a common reference genome using transanno (https://github.com/informationsea/transanno; accessed July 2026) to generate chain files (site-matching files) between the five reference genomes and to liftover the coordinate of all VCFs to a common reference genome G2_CRI. In addition, we used bcftools to correct errors introduced during coordinate transfer by first splitting multiallelic sites into separate records (“norm -m -any --force”) and removing duplicate records (“norm -d exact”). Finally, all coordinate-transferred VCFs were merged into one using bcftools (merge -m all), and only biallelic sites with a mean sequencing depth >5 and a missing rate <10% were retained (--min-alleles 2 --max-alleles 2 --remove-indels --max-missing 0.9 --mac 1 --minDP 5) using VCFtools. The final VCF was composed of 40 samples from each of the 16 species (exclude outgroup), representing all three genome groups.

Variant sites from the final VCF were first assessed for missing sites in each sample using VCFtools ("--missing"). The entire dataset was then filtered in PLINK v.1.96 to remove rare variants (“--mac 2”), prune variants in high linkage disequilibrium (“--indep-pairwise 50 10 0.2”), and exclude samples with more than 10% missing genotypes ("--mind 0.1"). Using the retained variants and samples, we assessed genetic relationships by principal component analysis (PCA) in PLINK (“--pca 20”) (Purcell et al., 2007). Population genetic structure was inferred using LEA (Frichot & François, 2015) by evaluating the number of ancestral populations (K) from 1 to 10, with 10 independent replicates performed for each K.

### 2.6 Introgression Detection from Genomic Data

Based on phylogenomic analysis and SNP data (see Results), we tested for alternative topologies among the outgroup *G. raimondii* (D5), and sections *Sturtia* (C genome), *Grandicalyx* (K genome), and *Hibiscoidea* (G genome), with the latter analyzed both as a single group and partitioned into two groups comprising G1 (*G. bickii*) and G2-G3 (*G. australe*–*G. nelsonii*). Individual samples were assigned to each group, and the relative weights of alternative topologies among these five groups were compared using Twisst or each previously generated, rerooted gene tree (Martin & Van Belleghem, 2017).

To investigate potential reticulate evolution, we inferred phylogenetic networks using SNaQ v1.2.0 (Solís-Lemus et al., 2017) with PhyloNetworks v1.4.0. Quartet concordance factors were calculated from all filtered and rerooted gene trees, with individuals assigned to their respective genetic groups (C1, C2, G1, G2–G3, K, and D5). Starting from the group-level ASTRAL topology, we inferred networks allowing a maximum of 0 to 3 reticulation events (‘hmax’), with 30 independent searches for each setting (‘runs’). The best network model was assessed using negative log-pseudolikelihood scores.

Genome-wide introgression among Australian cottons was assessed using the SNP-based *f*3 statistic implemented in ADMIXTOOLS v2.0.4 (Maier et al., 2023), which tests for excess allele sharing among all possible trios-species combinations using the previously filtered SNP dataset. In addition, Kmer-based interspecific introgression analyses using KmerCity (Garsmeur et al., 2025; Ning et al., 2026) was used to detect sequence overlap in highly repetitive genomic regions. Specifically, the trimmed reads were first filtered against a contaminant database (“Standard-8”) Kraken2 v.2.1.2 (Wood et al., 2019), followed by removal of organellar reads via mirabait (https://github.com/DrMicrobit/mira) using a customized database containing *G. hirsutum* mitochondrial and chloroplast DNA sequences (Ning et al., 2026). The resulting clean reads were randomly subsampled to 6 M (million) forward and reverse reads for downstream analyses. Given that most K group species were represented by a single sample, we classified all 38 samples into six genetic groups representing *G. sturtianum* (C1), *G. robinsonii* (C2), *G. bickii* (G1), *G. australe* (G2), *G. nelsonii* (G3), and section *Grandicalyx* (K), and calculated the average Kmer occurrence within each group in R (R. Team, 2014). Specifically, Kmers that were either shared among all genetic groups, restricted to a single group, or had a total occurrence of fewer than 10 across groups were excluded. The remaining overlapping Kmer patterns between groups were plotted in ggplot2 (Wilkinson, 2011).

## 3. Results

### 3.1 Genome Sequencing

Genetic data were generated and retrieved for 42 individuals representing 16 of the 17 Australian *Gossypium* species, ranging from one to ten individuals per species (Table 1). The only species not included was *G. pilosum* (K4), for which both fresh and herbarium specimens were unavailable (Table 1). Most samples averaged greater than 45.7-fold coverage, based on previously reported genome sizes (Hendrix & Stewart, 2005), ranging from 0.3-to 104.6-fold coverage, i.e., ∼4 million (M) to 1.66 billion (B) reads in samples *G. australe* (G2_SRR8666611) and *G. enthyle* (K7_Kew47098), respectively. All newly sequenced samples and most previously sequenced samples included more than 100 M reads, aside from two downloaded from NCBI, i.e., *G. robinsonii* (C2_robinsonii_SRR8666610) and *G. australe* (G2_australe_SRR8666611), which included only ∼4 M reads each (Table S1).

Because of the length of time involved in diversification of the Australian *Gossypium* clade (*circa* 5 – 10 million years (Cai et al., 2020; Cronn et al., 2002; Wendel & Albert, 1992)), we first assessed the potential for interspecific reference biases to distort mapping efficiency. Reads from each individual were mapped independently to the four available Australian *Gossypium* reference genomes: *G. sturtianum* (C1_HAU), *G. bickii* (G1_HAU), *G. australe* (G2_CRI), and *G. rotundifolium* (K2_HAU). Comparisons among mapping rates indicated that although the proportion of reads mapping to each reference (i.e., the primary alignment rates) remained high (91.71% - 99.52%) in all sample-versus-reference combinations, restricting the data to the proportion of reads pairs that mapped as expected pairs (i.e., the paired mapping rates) revealed more variation, typically yielding overall lower rates ranging from 47.58% – 96.34% (Fig. 3).

**Fig. 3.**
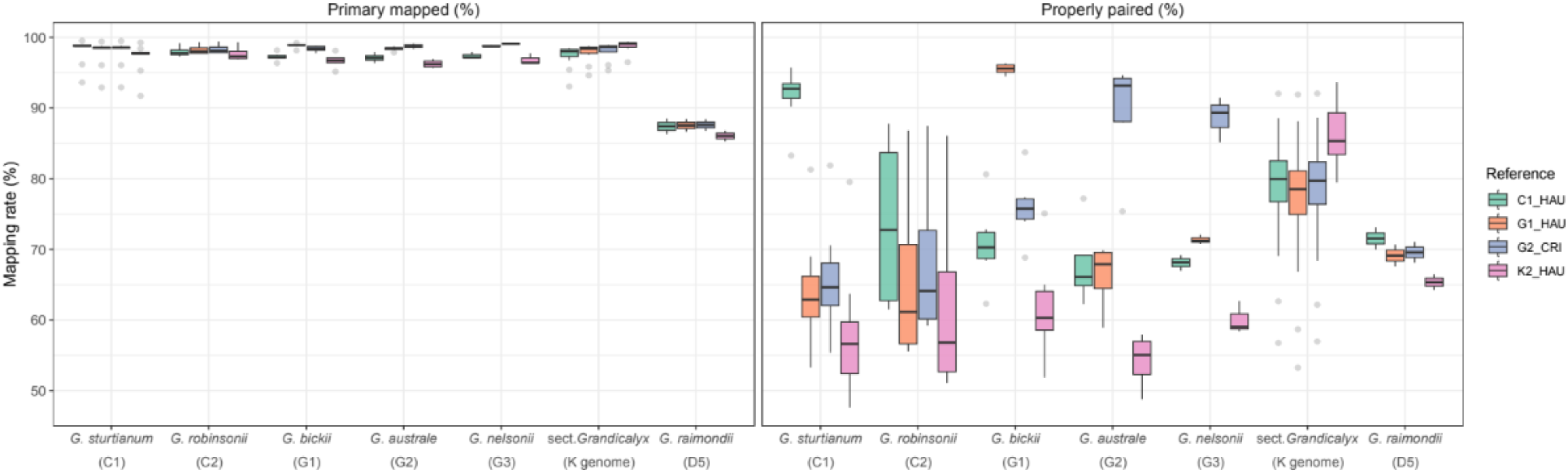
Mapping percentage comparisons for primary mapped reads (left) and paired-end mapped reads (right). The x-axis shows seven genome groups *Gossypium sturtianum* (C1), *G. robinsonii* (C2), *G. bickii* (G1), *G. australe* (G2), *G. nelsonii* (G3), section *Grandicalyx* (K genome), and the outgroup *G. raimondii* (D5) and the y-axis shows the mapping rate (%) against four reference genomes, with boxplots colored by reference genome.

Among the three taxonomic sections, the strongest reference biases were observed in *Sturtia* and *Hibiscoidea*, where the average proportion of reads mapped as proper pairs ranged from 89.1% to 95.5% when mapping intraspecifically but only averaged 65.0% when mapped interspecifically (Table S1; Fig. 3). In contrast, section *Grandicalyx* (K genome) exhibited relatively limited interspecific reference bias, consistent with the relatively recent radiation among its constituent species (Seelanan et al., 1999). Among the four Australian cotton reference genomes, the K2_HAU reference yielded the highest paired-mapping rate for the K-genome species, ranging from 79.5% to 93.6%. Lacking a species-specific reference, *G. robinsonii* (C2) and *G. nelsonii* (G3) showed the highest interspecific paired mapping rate when mapped to the reference in the same section, with *G. robinsonii* reaching 73.7% against the C1_HAU reference and *G. nelsonii* showing 71.2% and 88.7% against the G1_HAU and G2_CRI references, respectively. These interspecific and intersectional mapping rate differences reflect the level of genome divergence among Australian *Gossypium*, and reflect, at least in part, their collective differences in genome size (Table 1).

Previous research has shown that genome sizes within sections *Sturtia* and *Hibiscoidea* (i.e., C and G genome) are approximately twice the size of that in *G. raimondii* (D genome), and that those in section *Grandicalyx* (K genome) are nearly three times larger, largely due to lineage-specific transposable element dynamics (Hawkins et al., 2008; Hu et al., 2021; Huang et al., 2024; M. Wang et al., 2022). Thus, these differences in paired mapping efficiency are likely attributable to genomic differences resulting in large part from transposable-element activity combined along with evolutionary divergence. Congruent with this observation, the outgroup, small-genome species *G. raimondii* (D5) exhibited consistently lower mapping rates for all reference genomes, both in the number of primary- and paired-mapped reads, as expected from its greater evolutionary distance from Australian *Gossypium* (Wendel & Grover, 2015). To minimize analytical distortions that might arise from mapping bias, for the remainder of the analyses we used the closest available reference genome (see methods).

### 3.2 Identification of Single-Copy Orthologous Genes

To facilitate phylogenetic analyses using nuclear genes, we first identified a set of single copy orthologous genes among all sampled *Gossypium* species, including the phylogenetic outgroup *G. raimondii*. As detailed in the methods, we identified 39,353 orthogroups, including 19,029 that were inferred to be single-copy orthogroups. Synteny analyses showed that the majority of orthologous genes were retained in collinear positions (Fig. 4), as in (M. Wang et al., 2022), indicating strong conservation of chromosome synteny despite the differences in genome size and the divergence time between the Australian cottons and the outgroup *G. raimondii*, estimated to be 5 to 10 million years (Grover et al., 2015). Notably, and as in earlier reports (Sheng et al., 2023; M. Wang et al., 2022), synteny analysis between the four Australian reference genomes and the outgroup *G. raimondii* revealed a reciprocal translocation between chromosomes 05 and 13. Specifically, 1,922 loci from *G. raimondii* chromosome D05 matched chromosome 13 in Australian cottons, while approximately 329 loci from D13 matched chromosome 05 (Fig. 4).

**Fig. 4.**
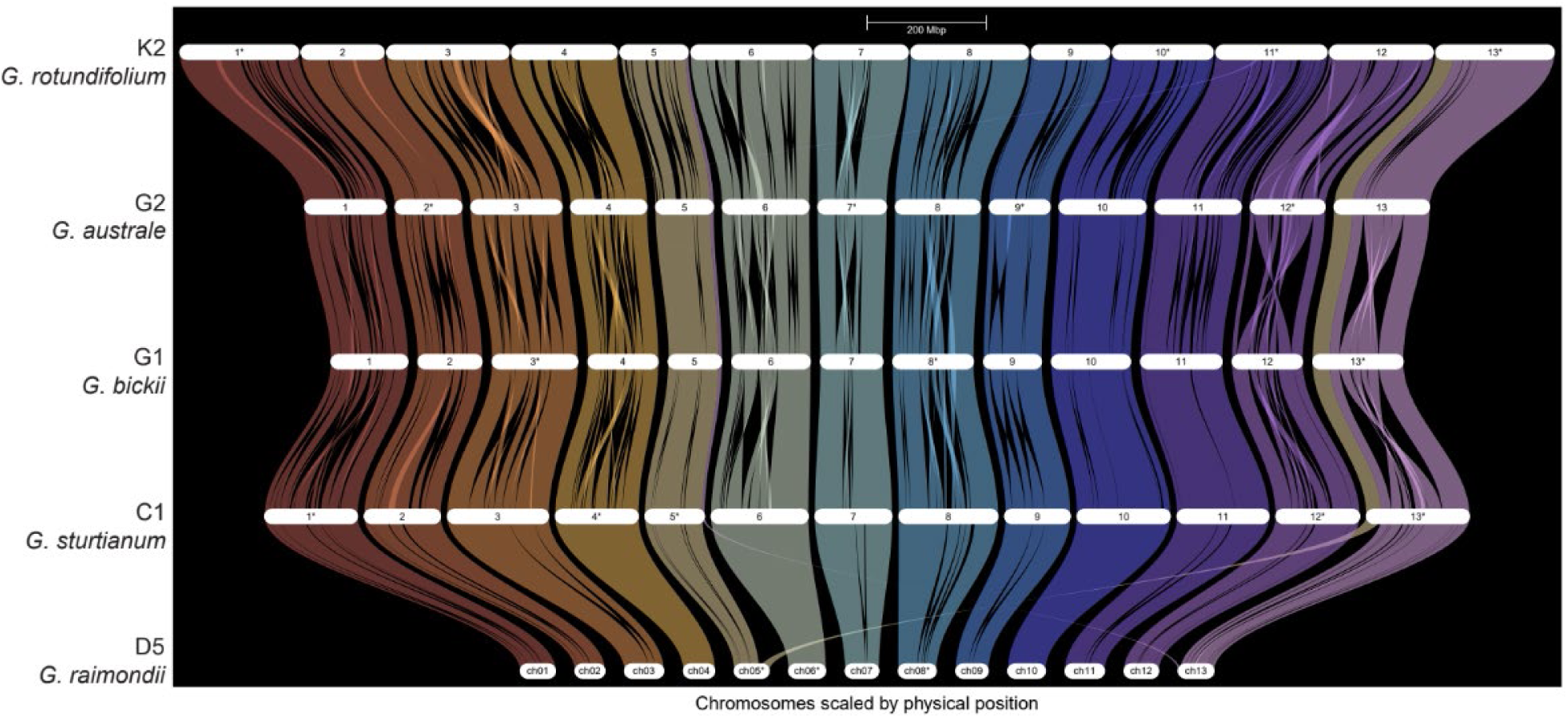
Synteny among five cotton reference genomes based on 39,353 orthogroups. Chromosomes (n=13) are positioned along the x-axis and are scaled by physical length.Syntenic orthologs are plotted according to their physical coordinates and connected across genomes on the y-axis by chromosome-colored lines, highlighting conserved chromosomal structure and structural rearrangements. Although largely syntenic, several inversions are detected (i.e., C1_HAU, *G. sturtianum*; G1_HAU, *G. bickii*; G2_CRI, *G. australe*; and K2_HAU, *G. rotundifolium*) in addition to a previously reported translocation between the outgroup *G. raimondii* (D5_T2t_PKU) and the Australian *Gossypium*.

### 3.3 Phylogenomic Analysis

Because phylogenetic relationships among Australian *Gossypium* species were previously characterized using a small number of nuclear markers (Seelanan et al., 1999; Wendel et al., 1991), we reassessed relationships among all 16 species using the rich dataset provided by 19,029 single-copy orthologs identified above (Fig. 4 Table S1). We removed loci that: (1) lacked data for all accessions (n = 224); (2) lacked parsimony-informative sites (n = 2); and (3) lacked phylogenetic resolution between Australian *Gossypium* species and the outgroup (n = 19), retaining 18,784 genes with a mean trimmed length of 2,481 bp and an average of 160 parsimony-informative sites per locus (Fig. S1). These data thus represent a rich set of genome-wide genic markers for phylogenetic reconstructions.

Phylogenetic analysis using ASTRAL recovered three monophyletic groups among Australian *Gossypium* species, corresponding to the previously recognized sections *Sturtia* (C genome), *Hibiscoidea* (G genome), and *Grandicalyx* (K genome) (Fig. 5a) (Craven et al., 1994; Liu et al., 2001; Seelanan et al., 1999; Stewart et al., 2015; Wendel et al., 1991). The more comprehensive data generated here provided stronger support for a closer relationship between the G and K genome species as a clade sister to the C genome species. Although the backbone of the phylogeny exhibited strong support, with 1.0 local posterior probabilities at nearly all internodes, the branch uniting the G and K genomes was relatively short, which is consistent with rapid radiation (Fig. 5a; Fig. S2). Divergence of sect. *Sturtia* from the common ancestor of sects. *Hibiscoidea* and *Grandicalyx* was estimated at 6.91 Ma, followed by the split between *Hibiscoidea* and *Grandicalyx* at 6.62 Ma. In addition, this latter node showed low concordance among gene trees, with only 19% supporting the inferred species-tree relationship, suggesting a rapid radiation of these lineages and hence limited phylogenetic support.

**Fig. 5.**
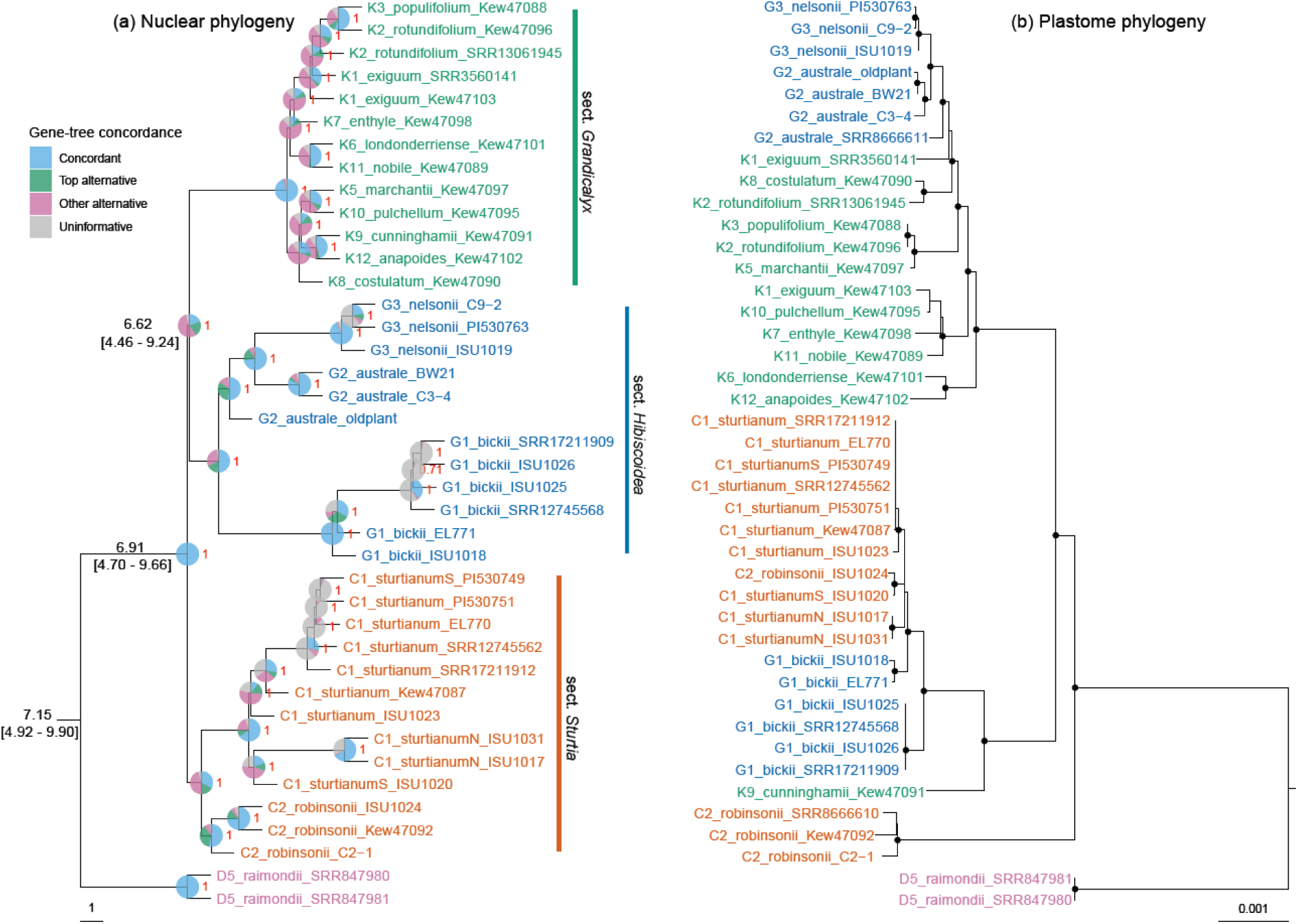
Phylogenetic inference of Australian *Gossypium* species. (a) Nuclear phylogeny inferred from 18,784 single-copy gene sequences. Terminal branch lengths were artificially set to 1. Tip labels are colored by three taxonomic sections: *Sturtia* (C genome; C1 and C2 in orange), *Hibiscoidea* (G genome; G1, G2, and G3 in blue), and *Grandicalyx* (K genome; K1 – K12 in green). Nodes are annotated with local posterior probabilities (0 – 1) and pie charts indicating gene-tree concordance with the species tree. The backbone of the phylogeny is annotated with sectional divergence times (Ma), with 95% highest posterior density intervals shown in brackets. (b) Plastome phylogeny. Nodes with bootstrap support of 100 are marked with black dots.

Within sect. *Sturtia* (C genome), two species, *G. sturtianum* (C1) and *G. robinsonii* (C2), formed two monophyletic clades; however, at the varietal level, the four individuals of *G. sturtianum* var. *nandewarense* (C1_sturtianumN) and *G. sturtianum* var. *sturtianum* (C1_sturtianumS) were recovered as paraphyletic, indicating that the evolutionary relationships among these varieties remains unresolved. Within sect. *Hibiscoidea* (G genome), a monophyletic lineage *G. nelsonii* (G3) was nested within *G. australe* (G2), and together they formed a clade sister to the monophyletic *G. bickii* (G1) lineage, consistent with previous studies (Seelanan et al., 1999; Wendel et al., 1991). Within sect. *Grandicalyx* (K genome), although most species were represented by only one accession, gene trees (95% concordance) strongly supported a split between two major clades, differing from the largely unresolved single nuclear marker ITS-based topology reported by Seelanan et al. (1999). The first clade included *G. costulatum* (K8), *G. anapoides* (K12), *G. cunninghamii* (K9), *G. pulchellum* (K10), and *G. marchantii* (K5). The second clade comprised *G. nobile* (K11), *G. londonderriense* (K6), *G. enthyle* (K7), *G. exiguum* (K1), *G. rotundifolium* (K2), and *G. populifolium* (K3). Despite this fundamental division within sect *Grandicalyx*, there do not appear to be any geographical correlates to explain this observation (Fig. S3). *Gossypium exiguum* (K1) and *G. rotundifolium* (K2) each included two accessions that were not recovered as monophyletic, suggesting that more extensive population-level sampling is required to resolve their intraspecific variation and interspecific relationships.

### 3.4 Plastome Phylogeny

The whole-plastome trimmed alignment (excluding one of the two inverted repeats, IRB) included 42 accessions, each with 132 kbp of cpDNA sequence, with 1,927 parsimony-informative sites. In contrast, the concatenated sequence of 84 plastid-encoded genes was 84 kbp in length and included less than half (825) as many parsimony-informative sites. Phylogenetic analysis of the whole-plastome sequences exhibited a different topology than that inferred from the nuclear genome dataset described above (Fig. 5a, 5b), despite generally high bootstrap support on the backbone of the phylogeny. Most notably, all three monophyletic sections in the nuclear-based phylogeny exhibited paraphyly in the plastome tree. Specifically, three of the four *G. robinsonii* (C2) accessions formed the earliest diverged lineage in the plastome phylogeny, and its sister node diverged into two major clades: (1) one containing *G. cunninghamii* (K9), all samples of *G. bickii* (G1) and *G. sturtianum* (C1), and the single remaining *G. robinsonii* (C2) sample [topology: (K9, (G1, C1–C2))]; and (2) the other containing the remaining species from sections *Grandicalyx* and *Hibiscoidea* [topology: (K1-3, 5-8, 10-12, (G2, G3))]. Notably, K-genome species (sect. *Grandicalyx*) represented the earliest diverging lineage in both clades; however, interspecific relationships within *Grandicalyx* (K genome) were not stable between the nuclear and cpDNA trees. For both *G. exiguum* (K1) and *G. rotundifolium* (K2), the two accessions representing each species were split in both the nuclear and cpDNA trees, highlighting their recent divergence and the possibility of lineage sorting (Lee-Yaw et al., 2019).

The presence of two multi-section clades suggests a history of introgression captured in the matrilinearly inherited chloroplast genome. These results largely recapitulated patterns observed in previous studies (Seelanan et al., 1999; Wendel et al., 1991), in which representatives of all three sections, i.e., *G. cunninghamii* (K9), *G. bickii* (G1), and *G. sturtianum* (C1), shared a similar chloroplast haplotype. Notably, only one of the *G. robinsonii* accessions (i.e., C2_robinsonii_ISU1024) was nested in the C1 clade, indicating possible historical introgression between *G. sturtianum* (C1) and *G. robinsonii* (C2). We also note that phylogenetic reconstructions using only 825 parsimony-informative sites from cpDNA coding genes revealed the same species relationships (Fig. S4), indicating the stability of these inferred relationships.

### 3.5 SNP-Based Genetic Relationships

Given the clear conflict between nuclear and cpDNA trees, and the reticulation suggested by cpDNA alone, we used several SNP-based approaches to test for historical, interspecific hybridization. Despite the conservation of gene content and synteny among protein-coding genes (Fig. 4), extensive genomic divergence driven by repeat element expansion (M. Wang et al., 2022) hindered using a single reference genome for SNP calling. Using the closest representative genome sequence for mapping each sample (Table S2), and a liftover-style method to place SNPs onto a common reference genome (G2_CRI), only 8% to 27.2% of interspecific genomic positions were retained as shared sites that could be jointly genotyped across species. Using this method, we identified 3.3 M high-quality biallelic variant sites across 36 individuals, with <10% missing data per site and <20% missing data per sample (Fig. 6a), which were LD-pruned to 227,379 biallelic variant sites for downstream analyses.

**Fig 6.**
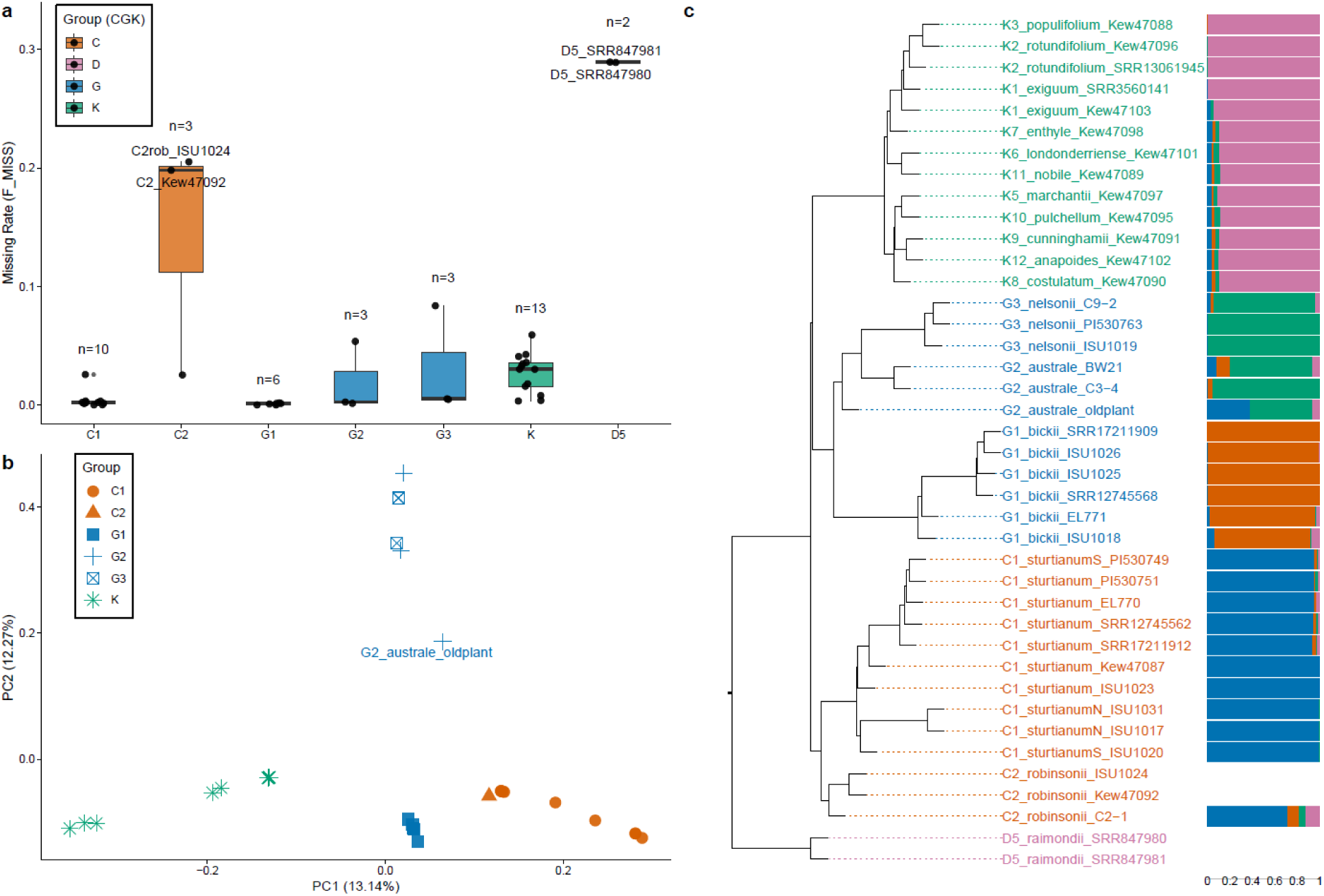
Genotyping missing rate, PCA, and LEA genetic structure analysis of Australian *Gossypium* samples. (a) Distribution of genotype missing rates among samples. (b) PCA showing genetic relationships among sections *Sturtia* (C genome; C1 and C2), *Hibiscoidea* (G genome; G1, G2, and G3), and *Grandicalyx* (K genome; K1 – K12), with colors representing genome groups and shapes representing species within the C and G genomes. (c) ASTRAL species tree and LEA genetic structure analysis (C2_robinsonii_ISU1024 and C2_robinsonii_Kew47092 were excluded due to high genotype missingness) identifying four major ancestral components.

We first used these SNPs to generate a multidimensional perspective on allelic content using Principal Component Analysis (PCA; Fig. 6b). Samples from each section were generally clustered, with the sections clearly separated along PC1 (13.14%). Species from sections *Sturtia* (C genome) and *Grandicalyx* (K genome) were placed on either end of PC1 while the species from section *Hibiscoidea* (G genome) were intermediate. PC2 (12.27%) generally separated *G. australe* (G2) and *G. nelsonii* (G3) from *G. bickii* (G1), where *G. bickii* was in between sections *Sturtia* and *Grandicalyx* on the first principal component but isolated from the other two species in sect. *Hibiscoidea* along PC2.

Genetic structure analysis recapitulated the PCA, identifying 4 groups as the optimal number of ancestral populations (K = 4) (Fig. 6c; Fig. S5). Samples were generally separated by section, albeit with sect. *Hibiscoidea* (G genome) separated into two groups: *G. bickii* (G1) versus *G. australe* (G2) and *G. nelsonii* (G3). No clear admixture patterns were observed, except for one individual (G2_australe_oldplant), which showed >20% ancestry from the sect. *Sturtia* (C genome); however, the timing of this introgression is unknown, possibly resulting from historical inadvertent hybridization in germplasm collections.

### 3.6 Reticulate Evolutionary History

Given the short internode separating the three sections (Fig. 5a), we tested alternative topologies among 18,784 gene trees for five well-defined groups (Fig. 5a; Fig. 6c) using Twisst. We analyzed four groups at a time, treating sect. *Hibiscoidea* either as a single group or as separate *G. bickii* (G1) and *G. australe–G. nelsonii* (G2–G3) lineages, consistent with the phylogeny (Fig. 5). The alternative topologies showed approximately equal representation of gene trees supporting sister relationships of sect. *Hibiscoidea* (G genome) with either sect. *Sturtia* (C genome; T1 36.1% in Fig. 7a) or sect. *Grandicalyx* (K genome; T3 38.5%). Similarly, this pattern was recapitulated for the group *G. australe–G. nelsonii* (G2–G3), for which the corresponding topologies received 38.6% and 37.7% support, respectively. In contrast, *G. bickii* (G1) showed stronger support (T3 40%) for being sister to sect. *Grandicalyx* (K genome) than with to sect. *Sturtia* (33.9%, C genome; T1). Notably, the topology with sections *Sturtia* (C genome) and *Grandicalyx* (K genome) as sister to each other received the lowest representation across all comparisons (T2 topology received average 25.23% supporting gene trees). These patterns are consistent with the PCA analysis and genetic structure (Fig. 6a, 6b), in indicating some level of “intermediacy” for *G. bickii* (G1) and the sister species *G. australe* and *G. nelsonii* (G2–G3) within sect. *Hibiscoidea*. Admixture *f*3 tests across all possible trios of Australian cotton species ranged from -0.0178 to 0.2862, with only four trios showing Z scores < -3, all associated with the sister species *G. australe* and *G. nelsonii* (G2–G3) and none providing significant evidence of intersectional introgression (Fig. S6a).

**Fig 7.**
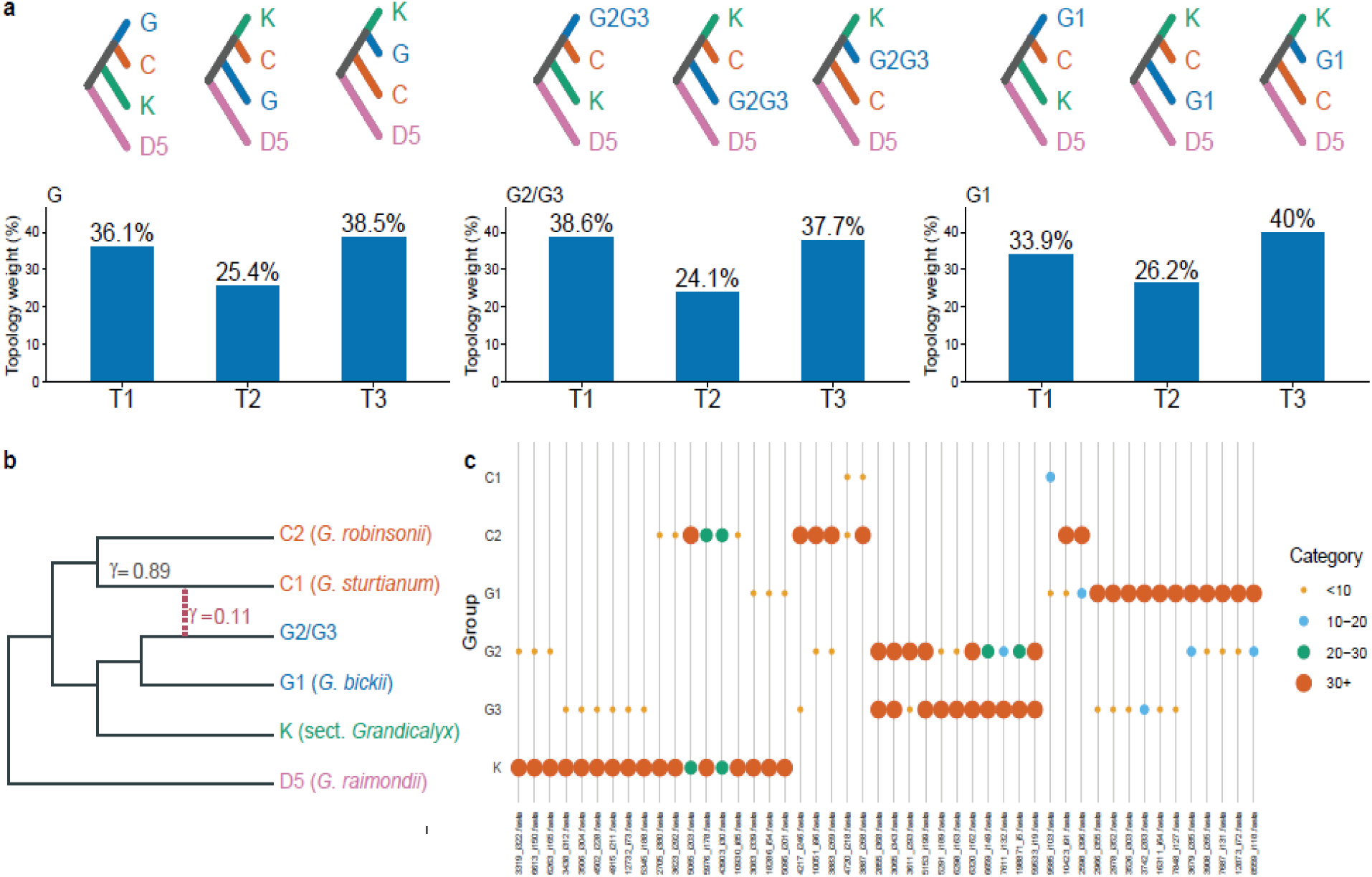
Inference of interspecific reticulation among Australian cottons. (a) Twisst analysis showing the three alternative topologies (top) for each comparison between four groups and the proportion of gene trees supporting each topology (bottom). (b) SNaQ-inferred network among six cotton groups, with the dashed arrow representing the inferred reticulation edge and γ indicating inheritance probabilities. (c) Kmer overlap analysis showing only Kmers shared between two groups. The complete *f*3 statistics and Kmer overlap analysis is provided in Fig. S5.

Quartet concordance-based network inference using SNaQ did not recover hybrid origins for *G. bickii* (G1) or *G. cunninghamii* (K9), but instead suggested reticulate ancestry for *G. sturtianum* (C1), involving the lineage leading to *G. robinsonii* (C2; with inheritance probabilities γ = 0.89) and the common ancestor of *G. australe* and *G. nelsonii* (G2–G3; γ = 0.11; Fig. 7c). In addition, we surveyed each sample for introgression using Kmer-overlapping patterns, as successfully used to identify introgression in sugarcane (Garsmeur et al., 2025). Kmer abundance patterns (Fig. 7b; Fig S5b) showed that 72,202 Kmers were uniquely shared between *Grandicalyx* (K genome) species and *G. robinsonii* (C2), many of which exhibited high abundance in both sections. An additional 79,721 K-mers were shared between sect. *Grandicalyx* (K genome) species and individual species of sect. *Hibiscoidea* (G genome) albeit typically at low frequencies within each sect. *Hibiscoidea* species (<10x). In addition, 305,267 distinct Kmers were shared exclusively between *G. australe* (G2) and *G. nelsonii* (G3) (Fig. S5b), consistent with their close phylogenetic relationship as a sister-species pair. However, the Kmer overlap patterns did not provide clear support for the proposed historical hybrid origins of *G. bickii* (G1), *G. cunninghamii* (K9), or *G. sturtianum* (C1) (Seelanan et al., 1999; Wendel et al., 1991).

## 4. Discussion

Australia has one of the highest levels of plant endemism globally, with approximately 84% of its native species restricted to the continent (Broadhurst & Coates, 2017). As wild relatives of the world’s most important cultivated cotton species, *Gossypium hirsutum* (Meyer & Dew, 2021; Ning et al., 2026; Ning, Rogers, et al., 2024), the Australian endemic *Gossypium* species comprise a distinct clade and represent one of the least explored lineages in the genus, despite exhibiting remarkable genomic and morphological diversity (Craven et al., 1994; Dubey et al., 2025; Hu et al., 2021). Here we reevaluate the origin and diversification of the Australian cotton lineage, using whole-genome resequencing data from 16 Australian *Gossypium* species, providing an evolutionary framework. Our specific goals were to provide a modern, whole-genome based perspective on inter- and intra-sectional relationships, and to evaluate whether there are nuclear data that support the potential hybrid or introgressive origin of *G. bickii* and *G. cunninghamii* (K9), as suggested by early molecular marker analyses (Seelanan et al., 1999; Wendel et al., 1991).

### 4.1 Origin and Diversification of Australian Cottons

Long-distance transoceanic dispersal is not uncommon in the genus *Gossypium* (Hu et al., 2021). All native Australian cotton species were previously suggested to descend from a single common ancestor that diverged from its diploid relatives in Africa and the Americas approximately 5 to 10 million years ago (Grover et al., 2015; Wendel & Albert, 1992). Our results confirmed a unique reciprocal chromosome translocation event involving chromosomes 5 and 13 that is shared by all three Australian cotton sections (Fig. 4), thus providing strong genetic evidence for a monophyletic origin of all Australian cotton species (Sheng et al., 2023; M. Wang et al., 2022). In addition, similar patterns of colonisation through long-distance dispersal followed by rapid diversification within Australia have been inferred for other plant lineages, including Camphorosmeae (Hühn et al., 2024), *Livistona* (Crisp et al., 2010), and native tobaccos (Cauz-Santos et al., 2024).

Upon arrival in Australia, the ancestral *Gossypium* lineage began to diversify. Our expanded dataset, derived from whole-genome resequencing data and comprising SNPs from over 18,000 orthologs across 40 accessions, unambiguously demonstrates that this diversification generated three monophyletic groups corresponding to the three taxonomic sections: *Sturtia* (C genome), *Hibiscoidea* (G genome), and *Grandicalyx* (K genome). Our results reveal particularly short internodes connecting the three sections (Fig. 5a), with a high-level of discordance between gene trees (Fig. 7a). These patterns are consistent with rapid diversification of the three clades, estimated here to have diverged *c*. 6 – 7 Ma (Fig. 5a), early in the radiation of Australian cottons and broadly coinciding with increasing aridification during the late Miocene, when the open and dry habitat expansion began (Byrne et al., 2008; Cauz-Santos et al., 2024; Crisp & Cook, 2013). The geographic origin of the ancestral Australian lineage remains inconclusive, however, given its position near the early divergence separating New World diploid cottons (subgenus *Houzingenia*) from the remaining *Gossypium* lineages (Hu et al., 2021; Wendel, 1989). Resolving its relationship to African and other diploid *Gossypium* lineages will require broader taxon sampling and additional genome-wide evidence.

### 4.2 Phylogenetic Relationships within Australian Cottons Sections

Extensive speciation and lineage-specific evolutionary diversification characterize the species-rich Kimberly region of NW Australia, which is recognized both for its megadiversity and microendemism (Pepper et al., 2014). This region is home to most species of sect. *Grandicalyx*, the only cotton species to evolve ant-mediated dispersal (Seelanan et al., 1999) (Fig. 2h). Rapid diversification within sect. *Grandicalyx* is estimated to have begun around 4 Ma, during the mid-Pliocene (Fig. S2). Alternating dry and wet climatic conditions in the Kimberley following this period may have promoted habitat fragmentation, leading to population isolation and divergence and ultimately contributing to speciation (Pepper et al., 2014; Stuut et al., 2019). In contrast, discordance between the nuclear and plastome phylogenies of taxa within sect. *Grandicalyx* (e.g., *G. exiguum* and *G. rotundifolium*; Fig. 5), suggests potential interspecific gene flow within the Kimberley region, although broader population-level sampling is needed to evaluate the extent. Such gene flow among rapidly diverging species may have blurred species boundaries, as documented in *Seringia* in the Kimberley region (Binks et al., 2020). Interesting among these is *G. cunninghamii* (K9), the only species in sect. *Grandicalyx* occurring outside of the Kimberley region, whose distribution is restricted to the Cobourg Peninsula and Kakadu National Park (Fig. 1). This species is nested within a well-defined nuclear clade of sect. *Grandicalyx* (Fig. 5) comprising *G. costulatum* (K8), *G. anapoides* (K12), *G. pulchellum* (K10), and *G. marchantii* (K5), yet it harbours a distinct cpDNA lineage sister to those of all other species in the section (Seelanan et al., 1999), potentially reflecting its geographic isolation and distinct evolutionary history (discussed below).

With respect to three species in sect. *Hibiscoidea*, *G. bickii* was recovered as a well-supported monophyletic lineage that diverged from the remainder of the section around 5.45 Ma (Fig. S2). *Gossypium bickii* is sister to but more distant from the species pair *G. australe* and *G. nelsonii* (Fig. 5; Fig. 6c), as expected based on earlier phylogenetic work (Seelanan et al., 1999; Stewart et al., 1987; Wendel et al., 1991). Although all three species have overlapping distributions in the central arid region of Australia, the divergence between *G. bickii* and the *G. australe–G. nelsonii* lineage suggests distinct evolutionary histories and possibly different ancestral origins. In addition, the closely related species pair *G. australe* and *G. nelsonii* were distinct, as demonstrated in earlier studies (Wendel et al., 1991; Wendel & Albert, 1992), but the more narrowly distributed *G. nelsonii* appears nested within the broadly distributed *G. australe* in both the nuclear and plastome phylogenies (Fig. 5). Traditional diagnostic morphological characters include vegetative pubescence, calyx morphology, foliar nectary position, and capsule surface characteristics (Stewart et al., 1987), some of which appear variable upon specimen examination (J. F. Wendel, pers. comm.), and may suggest interspecific gene flow in areas of contact.

Further discordance between datasets was detected in the two largely allopatric species of sect. *Sturtia*. *Gossypium robinsonii* is restricted to Western Australia whereas *G. sturtianum* is more widely distributed across Australia’s arid interior. *Gossypium robinsonii* resolved as monophyletic in the nuclear reconstructions but polyphyletic in the cpDNA tree, due to one accession (ISU1024) bearing an apparent *G. sturtianum-*like chloroplast genome (Fig. 5b). The remaining three *G. robinsonii* accessions formed a distinct clade representing the earliest-diverging plastome lineage among the Australian cotton species sampled. Thus, the unexpected occurrence of a “foreign cytoplasm” similar to that of modern *G. sturtianum* is extended here to a total of three geographically widely disjunct species that belong to three different taxonomic sections that are otherwise each monophyletic, implicating a history of introgression between the three sections.

### 4.3 Phylogenetic Incongruent between Australian Cottons Sections

Results from the present study recapitulated the previously observed cytonuclear discordance (Seelanan et al., 1999; Wendel et al., 1991)(Fig. 5a), which early studies interpreted as evidence of interspecific reticulation. Indeed, the nuclear and plastome-based discordance for *G. bickii* and *G. cunninghamii* are reiterated here using expanded taxonomic sampling and comprehensive (whole-genome) molecular phylogenomic analyses (Fig. 5a). In the cpDNA tree, *G. bickii* and *G. cunninghamii* grouped with *G. sturtianum*, with *G. cunninghamii* occupying a more basal position. In contrast, the pair was nested within the other major plastome clade containing the remaining species in *Grandicalyx* (Fig. 5b). These contrasting placements suggest distinct maternal histories for *G. bickii* and the *G. australe*–*G. nelsonii* lineages. In addition, the intermediate position of sect. *Hibiscoidea* (G genome) along PC1 in the SNP-based PCA (Fig. 6b), together with the phylogenetic discordance, raises the possibility of ancestral genetic exchange involving sections *Sturtia* (C genome) and *Grandicalyx* (K genome).

Our genome-wide analyses detected no compelling evidence of nuclear introgression (Fig S4), except that the SNaQ network analysis suggested a possible hybrid origin of *G. sturtianum*, involving lineages related to *G. robinsonii* (C2) and the *G. australe–G. nelsonii* pair (G2–G3; Fig. 7b). This phylogenetic incongruence may nonetheless reflect one or more responsible evolutionary processes such as incomplete lineage sorting, ancient hybridization or introgression, and rapid radiation. The ability to identify genomic signatures of interspecific hybridization generally depends on the timing, extent, and frequency of gene flow (Baack & Rieseberg, 2007; Groh & Coop, 2024). Ancient hybridization events may leave weak or fragmented genomic signatures due to recombination, population demographic history, selection, and genetic drift, whereas recent hybridization events are more likely to retain more intact introgressed genomic segments that are more readily detected (Harrison & Larson, 2014; Payseur & Rieseberg, 2016; Racimo et al., 2015; Taylor & Larson, 2019). Alternatively, the observed cytonuclear discordance could reflect chloroplast “capture”, whereby a rare immigrant introduced the maternal cytoplasm, while repeated backcrossing with pollen from a much larger resident population progressively diluted its nuclear contribution, retaining the immigrant chloroplast genome but leaving little detectable nuclear evidence of introgression (Lee-Yaw et al., 2019; Rieseberg & Wendel, 1993). In addition, rapid species radiation may allow ancestral polymorphisms to persist across successive speciation events, generating phylogenetic discordance through incomplete lineage sorting (Joly et al., 2009; Lee-Yaw et al., 2019). Given the substantial divergence and relative antiquity of the cytonuclear discordance, it is unclear what if any data can be brought to critically bear on its genesis.

An alternative to the foregoing is one in which the ancestral lineage leading to *G. robinsonii* (C2) also diversified into section *Grandicalyx* (K genome), followed by ancient hybridization/introgression between these two lineages, leading to the the section *Hibiscoidea* (G genome) and *G. sturtianum*. It is conceivable that the initial colonization of the Australian continent occurred in the northwestern part of the continent, a proposal consistent with its relative proximity to related African *Gossypium* clades and from the phylogenetic reconstructions shown here. This initial colonization could subsequently spread across much of the arid (section *Hibiscoidea*) and seasonally arid (section *Grandicalyx*) continent, evolving a wide range of growth forms drought adaptations, reproductive biology and dispersal ecology (Craven et al., 1994; Dubey et al., 2025; Fryxell, 1992; Masoomi-Aladizgeh et al., 2022). A question that arises is whether ancient interspecific introgression may have contributed to this evolutionary radiation, perhaps via the transfer of beneficial alleles among diverging lineages, a process increasingly recognized as an important driver of adaptation in plants (Rieseberg & Wendel, 1993; Schmickl & Yant, 2021). Together with agriculturally valuable traits such as delayed gossypol synthesis in developing seeds (Brubaker et al., 1996; Cheng et al., 2021; Jan et al., 2022; Sheng et al., 2023), these characteristics make Australian cottons an exceptional system for studying genome evolution, environmental adaptation, and the genetic basis of traits relevant to cotton improvement.

## Author contributions

W.N., C.E.G., and J.F.W. co-designed the experimental analyses. W.N. wrote the first draft of the manuscript and created all figures. C.E.G. and J.F.W. supervised the research and revised the manuscript. W.N., C.E.G., M.A.A., and J.A.U. contributed to data generation and analysis. J.A.U., and J.F.W. acquired funding.

## Data availability

All relevant code and scripts for the analyses presented here are available through Github (https://github.com/Wendellab/CGKcotton). Sequence reads for greenhouse maintained samples are available under PRJNA539957, and sequence reads for herbarium samples are under NCBI.

## Competing of interests statement

The authors declare no conflict of interest.

## Acknowledgments

The authors would like to thank Emma Miller for growing and maintaining the greenhouse samples, and Justin Conover for assistance in collecting leaves for sequencing. The authors also thank the Royal Botanic Gardens, Kew for access to herbarium material, specifically Imalka Kawandahala and the Kew Herbarium staff for assistance in collecting some of the plant material used in this study. The authors used AI-assisted tools GPT-6–based Codex for language polishing and bioinformatics pipeline development. All AI-assisted content and code were reviewed and verified by W.N., who takes full responsibility for the final manuscript and analyses.

## Funding

The research reported in this paper is partially supported by the HPC@ISU equipment at Iowa State University, some of which has been purchased through funding provided by NSF under MRI grants number 1726447 and MRI2018594. Additional support was provided by USDA-ARS under awards 59-6066-4-006 (Genomics of Kenaf, Cotton, and Other Malvaceae; to MAA) and 59-6066-4-007 (Investigating Agricultural & Environmental Systems; to MAA). This work was funded through grants from NSF and Cotton Inc. to J.A.U. and J.F.W. Part of this work was supported through by the National Geographic Society to J.F.W..

## 6. Supplemental materials

Fig. S1. Distribution of trimmed alignment length (bp) and parsimony-informative sites across 18,784 loci.

Fig. S2. Bayesian divergence-time estimates for Australian cottons. Node labels indicate divergence times between lineages, blue bars show the 95% corresponding intervals.

Fig. S3. Geographic distributions of the two clades of sect. *Grandicalyx* in Western Australia. Colors and symbols distinguish species.

Fig. S4. Phylogeny inferred from chloroplast protein-coding genes, with bootstrap support values shown at internal nodes.

Fig S5. Cross-entropy values and the optimal number of ancestral populations.

Fig S6. Genetic *f*3 admixture analysis (a) and KmerCity-based (b) genomic composition analysis of k-mer overlap patterns.

Table S1. Sampling and sequencing summary for Australian *Gossypium* accessions. The table lists each accession’s species, genetic group, source, read count (millions), read length (bp), and estimated sequencing depth. Primary and paired-read mapping rates (%) are shown for four reference genomes: C1 (*G. sturtianum*), G1 (*G. bickii*), G2 (*G. australe*), and K2 (*G. rotundifolium*). The table identifies accessions used for nuclear and plastome analyses, indicates whether plastomes were assembled de novo or by reference mapping, and provides NCBI accession numbers.

Table S2. Genotyping sites statistics for 40 samples mapped to five reference genomes, and liftover to the G2 (*G. australe*) reference genome. Total genotyped sites (to assigned reference genomes), mapped to G2_GRI, unmapped to G2_GRI, and final sites after duplicate removal are shown.

## Notes

### Competing Interest Statement

The authors have declared no competing interest.

## References Cited

Abbott, R. J. (2017). Plant speciation across environmental gradients and the occurrence and nature of hybrid zones: Hybrid zones and plant speciation. Journal of Systematics and Evolution, 55(4), 238–258. 10.1111/jse.12267

Abbott, R. J., Barton, N. H., & Good, J. M. (2016). Genomics of hybridization and its evolutionary consequences. Molecular Ecology, 25(11), 2325–2332. 10.1111/mec.13685

Alvarez, I., & Wendel, J. F. (2003). Ribosomal ITS sequences and plant phylogenetic inference. Molecular Phylogenetics and Evolution, 29(3), 417–434. 10.1016/s1055-7903(03)00208-2

Baack, E. J., & Rieseberg, L. H. (2007). A genomic view of introgression and hybrid speciation. Current Opinion in Genetics & Development, 17(6), 513–518. 10.1016/j.gde.2007.09.001

Binks, R. M., Wilkins, C. F., Markey, A. S., Lyons, M. N., & Byrne, M. (2020). Genomic data and morphological re-assessment reveals synonymy and hybridisation among *Seringia* taxa (Lasiopetaleae, Malvaceae) in remote north-western Australia. Taxon, 69(2), 307–320. 10.1002/tax.12233

Bolger, A. M., Lohse, M., & Usadel, B. (2014). Trimmomatic: a flexible trimmer for Illumina sequence data. Bioinformatics, 30(15), 2114–2120. 10.1093/bioinformatics/btu170

Borowiec, M. L. (2016). AMAS: a fast tool for alignment manipulation and computing of summary statistics. PeerJ, 4(e1660), e1660. 10.7717/peerj.1660

Broadhurst, L., & Coates, D. (2017). Plant conservation in Australia: Current directions and future challenges. Plant Diversity, 39(6), 348–356. 10.1016/j.pld.2017.09.005

Brubaker, C. L., Benson, C. G., Millet, C., & Leach, D. N. (1996). Occurrence of terpenoid aldehydes and lysigenous cavities in the “Glandless” seeds of Australian *Gossypium* species. Australian Journal of Botany, 44(5), 601–612. 10.1071/bt9960601

Byrne, M., Yeates, D. K., Joseph, L., Kearney, M., Bowler, J., Williams, M. A. J., Cooper, S., Donnellan, S. C., Keogh, J. S., Leys, R., Melville, J., Murphy, D. J., Porch, N., & Wyrwoll, K.-H. (2008). Birth of a biome: insights into the assembly and maintenance of the Australian arid zone biota. Molecular Ecology, 17(20), 4398–4417. 10.1111/j.1365-294X.2008.03899.x

Cai, Y., Cai, X., Wang, Q., Wang, P., Zhang, Y., Cai, C., Xu, Y., Wang, K., Zhou, Z., Wang, C., Geng, S., Li, B., Dong, Q., Hou, Y., Wang, H., Ai, P., Liu, Z., Yi, F., Sun, M., … Liu, F. (2020). Genome sequencing of the Australian wild diploid species *Gossypium australe* highlights disease resistance and delayed gland morphogenesis. Plant Biotechnology Journal, 18(3), 814–828. 10.1111/pbi.13249

Capella-Gutiérrez, S., Silla-Martínez, J. M., & Gabaldón, T. (2009). trimAl: a tool for automated alignment trimming in large-scale phylogenetic analyses. Bioinformatics, 25(15), 1972–1973. 10.1093/bioinformatics/btp348

Cauz-Santos, L. A., Samuel, R., Metschina, D., Christenhusz, M. J. M., Dodsworth, S., Dixon, K. W., Conran, J. G., Paun, O., & Chase, M. W. (2024). Recent speciation and adaptation to aridity in the ecologically diverse Pilbara region of Australia enabled the native tobaccos (*Nicotiana*; Solanaceae) to colonize all Australian deserts. Molecular Ecology, 33(18), e17498. 10.1111/mec.17498

Cheng, H., Feng, X., Zuo, D., Zhang, Y., Wang, Q., Lv, L., Wu, C., Li, S., Dai, Y., Qu, D., He, M., Liu, S., & Song, G. (2021). Gene expression correlation analysis reveals MYC-NAC regulatory network in cotton pigment gland development. International Journal of Molecular Sciences, 22(9), 5007. 10.3390/ijms22095007

Craven, L. A., Stewart, J. M. D., Brown, A. H. D., & Grace, J. P. (1994). The Australian wild species of *Gossypium*. Challenging the Future: Proceedings of the World Cotton Research Conference I, 278–281.

Crisp, M. D., & Cook, L. G. (2013). How was the Australian flora assembled over the last 65 million years? A molecular phylogenetic perspective. Annual Review of Ecology, Evolution, and Systematics, 44(1), 303–324. 10.1146/annurev-ecolsys-110512-135910

Crisp, M. D., Isagi, Y., Kato, Y., Cook, L. G., & Bowman, D. M. J. S. (2010). *Livistona* palms in Australia: ancient relics or opportunistic immigrants? Molecular Phylogenetics and Evolution, 54(2), 512–523. 10.1016/j.ympev.2009.09.020

Cronn, R. C., Small, R. L., Haselkorn, T., & Wendel, J. F. (2002). Rapid diversification of the cotton genus (*Gossypium*: Malvaceae) revealed by analysis of sixteen nuclear and chloroplast genes. American Journal of Botany, 89(4), 707–725. 10.3732/ajb.89.4.707

Cronn, R. C., & Wendel, J. F. (2004). Cryptic trysts, genomic mergers, and plant speciation: Research review. The New Phytologist, 161(1), 133–142. 10.1111/j.1469-8137.2004.00947.x

Danecek, P., Auton, A., Abecasis, G., Albers, C. A., Banks, E., DePristo, M. A., Handsaker, R. E., Lunter, G., Marth, G. T., Sherry, S. T., McVean, G., Durbin, R., & 1000 Genomes Project Analysis Group. (2011). The variant call format and VCFtools. Bioinformatics, 27(15), 2156–2158. 10.1093/bioinformatics/btr330

Danecek, P., Bonfield, J. K., Liddle, J., Marshall, J., Ohan, V., Pollard, M. O., Whitwham, A., Keane, T., McCarthy, S. A., Davies, R. M., & Li, H. (2021). Twelve years of SAMtools and BCFtools. GigaScience, 10(2). 10.1093/gigascience/giab008

Demotte, P., Panchaksaram, M., Kumarasinghe, H., Ly-Trong, N., dos Reis, M., & Minh, B. Q. (2025). IQ2MC: A new framework to infer phylogenetic time trees using IQ-TREE 3 and MCMCTree with mixture models. In EcoEvoRxiv. 10.32942/x2cd2x

Dubey, G., Phillips, A. L., Kemp, D. J., & Atwell, B. J. (2025). Physiological and structural traits contribute to thermotolerance in wild Australian cotton species. Annals of Botany, 135(3), 577–588. https://academic.oup.com/aob/article-abstract/135/3/577/7709961

Edelman, N. B., Frandsen, P. B., Miyagi, M., Clavijo, B., Davey, J., Dikow, R. B., García-Accinelli, G., Van Belleghem, S. M., Patterson, N., Neafsey, D. E., Challis, R., Kumar, S., Moreira, G. R. P., Salazar, C., Chouteau, M., Counterman, B. A., Papa, R., Blaxter, M., Reed, R. D., … Mallet, J. (2019). Genomic architecture and introgression shape a butterfly radiation. Science (New York, N.Y.), 366(6465), 594–599. 10.1126/science.aaw2090

Elgvin, T. O., Trier, C. N., Tørresen, O. K., Hagen, I. J., Lien, S., Nederbragt, A. J., Ravinet, M., Jensen, H., & Sætre, G.-P. (2017). The genomic mosaicism of hybrid speciation. Science Advances, 3(6), e1602996. 10.1126/sciadv.1602996

Emms, D. M., & Kelly, S. (2019). OrthoFinder: phylogenetic orthology inference for comparative genomics. Genome Biology, 20(1), 238. 10.1186/s13059-019-1832-y

Endrizzi, J. E., Turcotte, E. L., & Kohel, R. J. (1985). Genetics, cytology, and evolution of Gossypium. In Advances in Genetics (pp. 271–375). Elsevier. 10.1016/s0065-2660(08)60515-5

Falster, D., Gallagher, R., Wenk, E. H., Wright, I. J., Indiarto, D., Andrew, S. C., Baxter, C., Lawson, J., Allen, S., Fuchs, A., Monro, A., Kar, F., Adams, M. A., Ahrens, C. W., Alfonzetti, M., Angevin, T., Apgaua, D. M. G., Arndt, S., Atkin, O. K., … Ziemińska, K. (2021). AusTraits, a curated plant trait database for the Australian flora. Scientific Data, 8(1), 254. 10.1038/s41597-021-01006-6

Folk, R. A., Soltis, P. S., Soltis, D. E., & Guralnick, R. (2018). New prospects in the detection and comparative analysis of hybridization in the tree of life. American Journal of Botany, 105(3), 364–375. 10.1002/ajb2.1018

Frichot, E., & François, O. (2015). LEA: An R package for landscape and ecological association studies. Methods in Ecology and Evolution / British Ecological Society. 10.1111/2041-210X.12382

Fryxell, P. A. (1965). A revision of the Australian species of *Gossypium* with observations on the occurrence of *Thespesia* in Australia (Malvaceae). Australian Journal of Botany, 13(1), 71–102. 10.1071/bt9650071

Fryxell, P. A. (1979). How to access research remotely. cabdirect.org. https://www.cabdirect.org/cabdirect/abstract/19812607534

Fryxell, P. A. (1992). A revised taxonomic interpretation of *Gossypium* L. (Malvaceae). Rheedea: Official Journal of Indian Association for Angiosperm Taxonomy, 2(2), 108–165. 10.22244/rheedea.1992.02.02.04

Fryxell, P. A., Craven, L. A., & Stewart, J. M. (1992). A revision of *Gossypium* sect. *Grandicalyx* (Malvaceae), including the description of six new species. Systematic Botany, 17(1), 91–114. 10.2307/2419068

Garsmeur, O., Rio, S., Pompidor, N., Lipzen, A., Hervouet, C., Durand, T., Daum, C., Yoshinaga, Y., Butterfield, M., Sanchez, A., Piperidis, G., Lincoln, N., Hale, A., Hoarau, J. Y., Terajima, Y., Lakshmanan, P., Sacks, E., Sharma, S., Vitrac, M., … D’Hont, A. (2025). The genomic footprints of wild *Saccharum* species trace domestication, diversification, and modern breeding of sugarcane. Cell, 188(25), 7252–7266.e15. 10.1016/j.cell.2025.09.017

Groh, J. S., & Coop, G. (2024). The temporal and genomic scale of selection following hybridization. Proceedings of the National Academy of Sciences of the United States of America, 121(12), e2309168121. 10.1073/pnas.2309168121

Grover, C. E., Arick, M. A., Thrash, A., Sharbrough, J., Hu, G., Yuan, D., Snodgrass, S., Miller, E. R., Ramaraj, T., Peterson, D. G., Udall, J. A., & Wendel, J. F. (2022). Dual domestication, diversity, and differential introgression in Old World cotton diploids. Genome Biology and Evolution, 14(12), evac170. 10.1093/gbe/evac170

Grover, C. E., Gallagher, J. P., Jareczek, J. J., Page, J. T., Udall, J. A., Gore, M. A., & Wendel, J. F. (2015). Re-evaluating the phylogeny of allopolyploid *Gossypium* L. Molecular Phylogenetics and Evolution, 92, 45–52. 10.1016/j.ympev.2015.05.023

Harrison, R. G., & Larson, E. L. (2014). Hybridization, introgression, and the nature of species boundaries. The Journal of Heredity, 105 Suppl 1(S1), 795–809. 10.1093/jhered/esu033

Hawkins, J. S., Hu, G., Rapp, R. A., Grafenberg, J. L., & Wendel, J. F. (2008). Phylogenetic determination of the pace of transposable element proliferation in plants: copia and LINE-like elements in *Gossypium*. Genome, 51(1), 11–18. 10.1139/g07-099

Hendrix, B., & Stewart, J. M. (2005). Estimation of the nuclear DNA content of *Gossypium* species. Annals of Botany, 95(5), 789–797. 10.1093/aob/mci078

Herbarium, W. A. (1998). FloraBase—the Western Australian Flora. http://www.vifabio.de/iqfBio/detail/1533

Hibbins, M. S., & Hahn, M. W. (2022). Phylogenomic approaches to detecting and characterizing introgression. Genetics, 220(2), iyab173. 10.1093/genetics/iyab173

Hu, G., Grover, C. E., Jareczek, J., Yuan, D., Dong, Y., Miller, E., Conover, J. L., & Wendel, J. F. (2021). Evolution and diversity of the cotton genome. In Cotton Precision Breeding (pp. 25–78). Springer International Publishing. 10.1007/978-3-030-64504-5_2

Huang, G., Bao, Z., Feng, L., Zhai, J., Wendel, J. F., Cao, X., & Zhu, Y. (2024). A telomere-to-telomere cotton genome assembly reveals centromere evolution and a Mutator transposon-linked module regulating embryo development. Nature Genetics, 56(9), 1953–1963. 10.1038/s41588-024-01877-6

Hühn, P., McDonald, J., Shepherd, K. A., & Kadereit, G. (2024). Diversification of Camphorosmeae (Amaranthaceae s.l.) during the Miocene-Pliocene aridification of inland Australia. Perspectives in Plant Ecology, Evolution and Systematics, 64(125811), 125811. 10.1016/j.ppees.2024.125811

Jan, M., Liu, Z., Guo, C., Zhou, Y., & Sun, X. (2022). An overview of cotton gland development and its transcriptional regulation. International Journal of Molecular Sciences, 23(9), 4892. 10.3390/ijms23094892

Jin, J.-J., Yu, W.-B., Yang, J.-B., Song, Y., dePamphilis, C. W., Yi, T.-S., & Li, D.-Z. (2020). GetOrganelle: a fast and versatile toolkit for accurate de novo assembly of organelle genomes. Genome Biology, 21(1), 241. 10.1186/s13059-020-02154-5

Joly, S., McLenachan, P. A., & Lockhart, P. J. (2009). A statistical approach for distinguishing hybridization and incomplete lineage sorting. The American Naturalist, 174(2), E54–70. 10.1086/600082

Junier, T., & Zdobnov, E. M. (2010). The Newick utilities: high-throughput phylogenetic tree processing in the UNIX shell. Bioinformatics, 26(13), 1669–1670. 10.1093/bioinformatics/btq243

Katoh, K., & Standley, D. M. (2013). MAFFT multiple sequence alignment software version 7: improvements in performance and usability. Molecular Biology and Evolution, 30(4), 772– 780. 10.1093/molbev/mst010

Kendig, K. I., Baheti, S., Bockol, M. A., Drucker, T. M., Hart, S. N., Heldenbrand, J. R., Hernaez, M., Hudson, M. E., Kalmbach, M. T., Klee, E. W., Mattson, N. R., Ross, C. A., Taschuk, M., Wieben, E. D., Wiepert, M., Wildman, D. E., & Mainzer, L. S. (2019). Sentieon DNAseq variant calling workflow demonstrates strong computational performance and accuracy. Frontiers in Genetics, 10, 736. 10.3389/fgene.2019.00736

Lamichhaney, S., Han, F., Webster, M. T., Andersson, L., Grant, B. R., & Grant, P. R. (2018). Rapid hybrid speciation in Darwin’s finches. Science (New York, N.Y.), 359(6372), 224–228. 10.1126/science.aao4593

Lee-Yaw, J. A., Grassa, C. J., Joly, S., Andrew, R. L., & Rieseberg, L. H. (2019). An evaluation of alternative explanations for widespread cytonuclear discordance in annual sunflowers (*Helianthus*). The New Phytologist, 221(1), 515–526. 10.1111/nph.15386

Liu, Q., Brubaker, C. L., Green, A. G., Marshall, D. R., Sharp, P. J., & Singh, S. P. (2001). Evolution of the FAD2-1 fatty acid desaturase 5’ UTR intron and the molecular systematics of *Gossypium* (Malvaceae). American Journal of Botany, 88(1), 92–102. 10.2307/2657130

Lovell, J. T., Sreedasyam, A., Schranz, M. E., Wilson, M., Carlson, J. W., Harkess, A., Emms, D., Goodstein, D. M., & Schmutz, J. (2022). GENESPACE tracks regions of interest and gene copy number variation across multiple genomes. eLife, 11(e78526), e78526. 10.7554/eLife.78526

Maier, R., Flegontov, P., Flegontova, O., Işıldak, U., Changmai, P., & Reich, D. (2023). On the limits of fitting complex models of population history to *f*-statistics. eLife, 12(e85492), e85492. 10.7554/eLife.85492

Martin, S. H., & Van Belleghem, S. M. (2017). Exploring evolutionary relationships across the genome using topology weighting. Genetics, 206(1), 429–438. 10.1534/genetics.116.194720

Masoomi-Aladizgeh, F., Kamath, K. S., Haynes, P. A., & Atwell, B. J. (2022). Genome survey sequencing of wild cotton (*Gossypium robinsonii*) reveals insights into proteomic responses of pollen to extreme heat. Plant, Cell & Environment, 45(4), 1242–1256. 10.1111/pce.14268

McKain, M. R., Johnson, M. G., Uribe-Convers, S., Eaton, D., & Yang, Y. (2018). Practical considerations for plant phylogenomics. Applications in Plant Sciences, 6(3), e1038. 10.1002/aps3.1038

Meyer, L., & Dew, T. (2021). Approved by USDA’s World Agricultural Outlook Board Cotton and Wool Outlook*: June* 2024. ers.usda.gov. https://www.ers.usda.gov/sites/default/files/_laserfiche/outlooks/109336/CWS-24f.pdf?v=62787

Minh, B. Q., Trifinopoulos, J., Schrempf, D., Schmidt, H. A., & Lanfear, R. (2019). IQ-TREE version 2.0: tutorials and Manual Phylogenomic software by maximum likelihood. *URL* Http://Www.Iqtree.Org. http://www.iqtree.org/doc/iqtree-doc.pdf

Mirarab, S., Reaz, R., Bayzid, M. S., Zimmermann, T., Swenson, M. S., & Warnow, T. (2014). ASTRAL: genome-scale coalescent-based species tree estimation. Bioinformatics, 30(17), i541–8. 10.1093/bioinformatics/btu462

Neph, S., Kuehn, M. S., Reynolds, A. P., Haugen, E., Thurman, R. E., Johnson, A. K., Rynes, E., Maurano, M. T., Vierstra, J., Thomas, S., Sandstrom, R., Humbert, R., & Stamatoyannopoulos, J. A. (2012). BEDOPS: high-performance genomic feature operations. Bioinformatics, 28(14), 1919–1920. 10.1093/bioinformatics/bts277

Nieto Feliner, G., Álvarez, I., Fuertes-Aguilar, J., Heuertz, M., Marques, I., Moharrek, F., Piñeiro, R., Riina, R., Rosselló, J. A., Soltis, P. S., & Villa-Machío, I. (2017). Is homoploid hybrid speciation that rare? An empiricist’s view. Heredity, 118(6), 513–516. 10.1038/hdy.2017.7

Nieto Feliner, G., Casacuberta, J., & Wendel, J. F. (2020). Genomics of evolutionary novelty in hybrids and polyploids. Frontiers in Genetics, 11, 792. 10.3389/fgene.2020.00792

Ning, W., Arick, M. A., 2nd, Udall, J. A., Hsu, C.-Y., Abdala-Roberts, L., Solís-Rodríguez, U., Briones-May, Y., Magbanua, Z. V., Pechanova, O., Bustos-Segura, C., Clancy, M. V., Díaz-Cruz, S., Garnica-Cabrera, A., Mamin, M., Yu, J. Z., Turlings, T. C. J., Hu, G., Peterson, D. G., Grover, C. E., & Wendel, J. F. (2026). Genomic diversity and the domestication history of cotton (*Gossypium hirsutum*). Proceedings of the National Academy of Sciences of the United States of America, 123(21), e2607107123. 10.1073/pnas.2607107123

Ning, W., Meudt, H. M., & Tate, J. A. (2024). A roadmap of phylogenomic methods for studying polyploid plant genera. Applications in Plant Sciences. 10.1002/aps3.11580

Ning, W., Rogers, K. M., Hsu, C.-Y., Magbanua, Z. V., Pechanova, O., Arick, M. A., 2nd, Kayal, E., Hu, G., Peterson, D. G., Udall, J. A., Grover, C. E., & Wendel, J. F. (2024). Origin and diversity of the wild cottons (*Gossypium hirsutum*) of Mound Key, Florida. Scientific Reports, 14(1), 14046. 10.1038/s41598-024-64887-8

Page, J. T., Liechty, Z. S., Huynh, M. D., & Udall, J. A. (2014). BamBam: genome sequence analysis tools for biologists. BMC Research Notes, 7(1), 829. 10.1186/1756-0500-7-829

Payseur, B. A., & Rieseberg, L. H. (2016). A genomic perspective on hybridization and speciation. Molecular Ecology, 25(11), 2337–2360. 10.1111/mec.13557

Pepper, M., Scott Keogh, J., & Ebach, M. (2014). Biogeography of the Kimberley, Western Australia: a review of landscape evolution and biotic response in an ancient refugium. Journal of Biogeography, 41(8), 1443–1455. 10.1111/jbi.12324

Pertea, G., & Pertea, M. (2020). GFF utilities: GffRead and GffCompare. F1000Research, 9, 304. 10.12688/f1000research.23297.2

Purcell, S., Neale, B., Todd-Brown, K., Thomas, L., Ferreira, M. A. R., Bender, D., Maller, J., Sklar, P., de Bakker, P. I. W., Daly, M. J., & Sham, P. C. (2007). PLINK: a tool set for whole-genome association and population-based linkage analyses. The American Journal of Human Genetics, 81(3), 559–575. 10.1086/519795

R. Team. (2014). R: A language and environment for statistical computing. MSOR Connections, 1. https://apps.dtic.mil/sti/citations/AD1039033

Racimo, F., Sankararaman, S., Nielsen, R., & Huerta-Sánchez, E. (2015). Evidence for archaic adaptive introgression in humans. Nature Reviews. Genetics, 16(6), 359–371. 10.1038/nrg3936

Rieseberg, L. H., & Wendel, J. F. (1993). Introgression and its consequences. https://dr.lib.iastate.edu/bitstreams/86e8cd99-abdc-45c2-8eef-80cac3d3014c/download

Runemark, A., Vallejo-Marin, M., & Meier, J. I. (2019). Eukaryote hybrid genomes. PLoS Genetics, 15(11), e1008404. 10.1371/journal.pgen.1008404

Schmickl, R., & Yant, L. (2021). Adaptive introgression: how polyploidy reshapes gene flow landscapes. The New Phytologist, 230(2), 457–461. 10.1111/nph.17204

Schumer, M., Rosenthal, G. G., & Andolfatto, P. (2018). What do we mean when we talk about hybrid speciation? Heredity, 120(4), 379–382. 10.1038/s41437-017-0036-z

Seelanan, T., Brubaker, C. L., Stewart, J. M., Craven, L. A., & Wendel, J. F. (1999). Molecular systematics of Australian *Gossypium* section *Grandicalyx* (Malvaceae). Systematic Botany, 24(2), 183–208. 10.2307/2419548

Sheng, K., Sun, Y., Liu, M., Cao, Y., Han, Y., Li, C., Muhammad, U., Daud, M. K., Wang, W., Li, H., Samrana, S., Hui, Y., Zhu, S., Chen, J., & Zhao, T. (2023). A reference-grade genome assembly for *Gossypium bickii* and insights into its genome evolution and formation of pigment glands and gossypol. Plant Communications, 4(1), 100421. 10.1016/j.xplc.2022.100421

Shumate, A., & Salzberg, S. L. (2021). Liftoff: accurate mapping of gene annotations. Bioinformatics (Oxford, England), 37(12), 1639–1643. 10.1093/bioinformatics/btaa1016

Small, R. L., Cronn, R. C., & Wendel, J. F. (2004). Use of nuclear genes for phylogeny reconstruction in plants. Australian Systematic Botany, 17(2), 145–170. 10.1071/sb03015

Smith, S. A., Moore, M. J., Brown, J. W., & Yang, Y. (2015). Analysis of phylogenomic datasets reveals conflict, concordance, and gene duplications with examples from animals and plants. BMC Evolutionary Biology, 15, 150. 10.1186/s12862-015-0423-0

Solís-Lemus, C., Bastide, P., & Ané, C. (2017). PhyloNetworks: A Package for Phylogenetic Networks. Molecular Biology and Evolution, 34(12), 3292–3298. 10.1093/molbev/msx235

Soltis, P. S., & Soltis, D. E. (2009). The role of hybridization in plant speciation. Annual Review of Plant Biology, 60(1), 561–588. 10.1146/annurev.arplant.043008.092039

Stewart, J. M., Craven, L. A., Brubaker, C., & Wendel, J. F. (2015). *Gossypium anapoides* (Malvaceae), a new species from Western Australia. Novon : A Journal for Botanical Nomenclature, 23(4), 447–451. 10.3417/2007140

Stewart, J. M., Fryxell, P. A., & Craven, L. A. (1987). The recongnition and geograpic distribution of *Gossypium nelsonii* (Malvaceae). Brunonia, 10(2), 215–218. 10.1071/bru9870215

Stuut, J.-B. W., De Deckker, P., Saavedra-Pellitero, M., Bassinot, F., Drury, A. J., Walczak, M. H., Nagashima, K., & Murayama, M. (2019). A 5.3-million-year history of monsoonal precipitation in northwestern Australia. Geophysical Research Letters, 46(12), 6946–6954. 10.1029/2019gl083035

Suarez-Gonzalez, A., Lexer, C., & Cronk, Q. C. B. (2018). Adaptive introgression: a plant perspective. Biology Letters, 14(3), 20170688. 10.1098/rsbl.2017.0688

Taylor, S. A., & Larson, E. L. (2019). Insights from genomes into the evolutionary importance and prevalence of hybridization in nature. Nature Ecology & Evolution, 3(2), 170–177. 10.1038/s41559-018-0777-y

Tillich, M., Lehwark, P., Pellizzer, T., Ulbricht-Jones, E. S., Fischer, A., Bock, R., & Greiner, S. (2017). GeSeq - versatile and accurate annotation of organelle genomes. Nucleic Acids Research, 45(W1), W6–W11. 10.1093/nar/gkx391

Tricou, T., Tannier, E., & de Vienne, D. M. (2022). Ghost lineages can invalidate or even reverse findings regarding gene flow. PLoS Biology, 20(9), e3001776. 10.1371/journal.pbio.3001776

Wang, K., Wendel, J. F., & Hua, J. (2018). Designations for individual genomes and chromosomes in *Gossypium*. Journal of Cotton Research, 1(1). 10.1186/s42397-018-0002-1

Wang, M., Li, J., Qi, Z., Long, Y., Pei, L., Huang, X., Grover, C. E., Du, X., Xia, C., Wang, P., Liu, Z., You, J., Tian, X., Ma, Y., Wang, R., Chen, X., He, X., Fang, D. D., Sun, Y., … Zhang, X. (2022). Genomic innovation and regulatory rewiring during evolution of the cotton genus *Gossypium*. Nature Genetics, 54(12), 1959–1971. 10.1038/s41588-022-01237-2

Wang, M., Li, J., Wang, P., Liu, F., Liu, Z., Zhao, G., Xu, Z., Pei, L., Grover, C. E., Wendel, J. F., Wang, K., & Zhang, X. (2021). Comparative genome analyses highlight transposon-mediated genome expansion and the evolutionary architecture of 3D genomic folding in cotton. Molecular Biology and Evolution, 38(9), 3621–3636. 10.1093/molbev/msab128

Wang, Y., Tang, H., Debarry, J. D., Tan, X., Li, J., Wang, X., Lee, T.-H., Jin, H., Marler, B., Guo, H., Kissinger, J. C., & Paterson, A. H. (2012). MCScanX: a toolkit for detection and evolutionary analysis of gene synteny and collinearity. Nucleic Acids Research, 40(7), e49. 10.1093/nar/gkr1293

Wendel, J. F. (1989). New World tetraploid cottons contain Old World cytoplasm. Proceedings of the National Academy of Sciences of the United States of America, 86(11), 4132–4136. 10.1073/pnas.86.11.4132

Wendel, J. F., & Albert, V. A. (1992). Phylogenetics of the cotton genus (*Gossypium*): Character-state weighted parsimony analysis of chloroplast-DNA restriction site data and its systematic and biogeographic implications. Systematic Botany, 17(1), 115–143. 10.2307/2419069

Wendel, J. F., & Grover, C. E. (2015). Taxonomy and evolution of the cotton genus, Gossypium. In Cotton (pp. 25–44). American Society of Agronomy, Inc., Crop Science Society of America, Inc., and Soil Science Society of America, Inc. 10.2134/agronmonogr57.2013.0020

Wendel, J. F., Stewart, J. M., & Rettig, J. H. (1991). Molecular evidence for homoploid reticulate evolution among Australian species of *Gossypium*. Evolution; International Journal of Organic Evolution, 45(3), 694–711. 10.1111/j.1558-5646.1991.tb04339.x

Wilkinson, L. (2011). ggplot2: Elegant Graphics for Data Analysis by WICKHAM, H. Biometrics, 67(2), 678–679. 10.1111/j.1541-0420.2011.01616.x

Wong, T. K. F., Ly-Trong, N., Ren, H., Demotte, P., Baños, H., Roger, A. J., Susko, E., Bielow, C., De Maio, N., Goldman, N., Hahn, M. W., Dos Reis, M., Vinh, L. S., Huttley, G., Lanfear, R., & Minh, B. Q. (2026). IQ-TREE 3: phylogenomic inference software using complex evolutionary models. Molecular Biology and Evolution, 43(5), msag117. 10.1093/molbev/msag117

Wood, D. E., Lu, J., & Langmead, B. (2019). Improved metagenomic analysis with Kraken 2. Genome Biology, 20(1), 257. 10.1186/s13059-019-1891-0

Yu, G., Smith, D. K., Zhu, H., Guan, Y., & Lam, T. T.-Y. (2017). Ggtree: An r package for visualization and annotation of phylogenetic trees with their covariates and other associated data. Methods in Ecology and Evolution / British Ecological Society, 8(1), 28–36. 10.1111/2041-210x.12628

Yu, J., Jung, S., Cheng, C.-H., Ficklin, S. P., Lee, T., Zheng, P., Jones, D., Percy, R. G., & Main, D. (2014). CottonGen: a genomics, genetics and breeding database for cotton research. Nucleic Acids Research, 42(Database issue), D1229–36. 10.1093/nar/gkt1064

Yuan, D., Grover, C. E., Hu, G., Pan, M., Miller, E. R., Conover, J. L., Hunt, S. P., Udall, J. A., & Wendel, J. F. (2021). Parallel and intertwining threads of domestication in allopolyploid cotton. Advancement of Science, 8(10), 2003634. 10.1002/advs.202003634

