## Supplemental figures for "Revisiting homoploid hybrid speciation in Australian cotton (*Gossypium*) species"

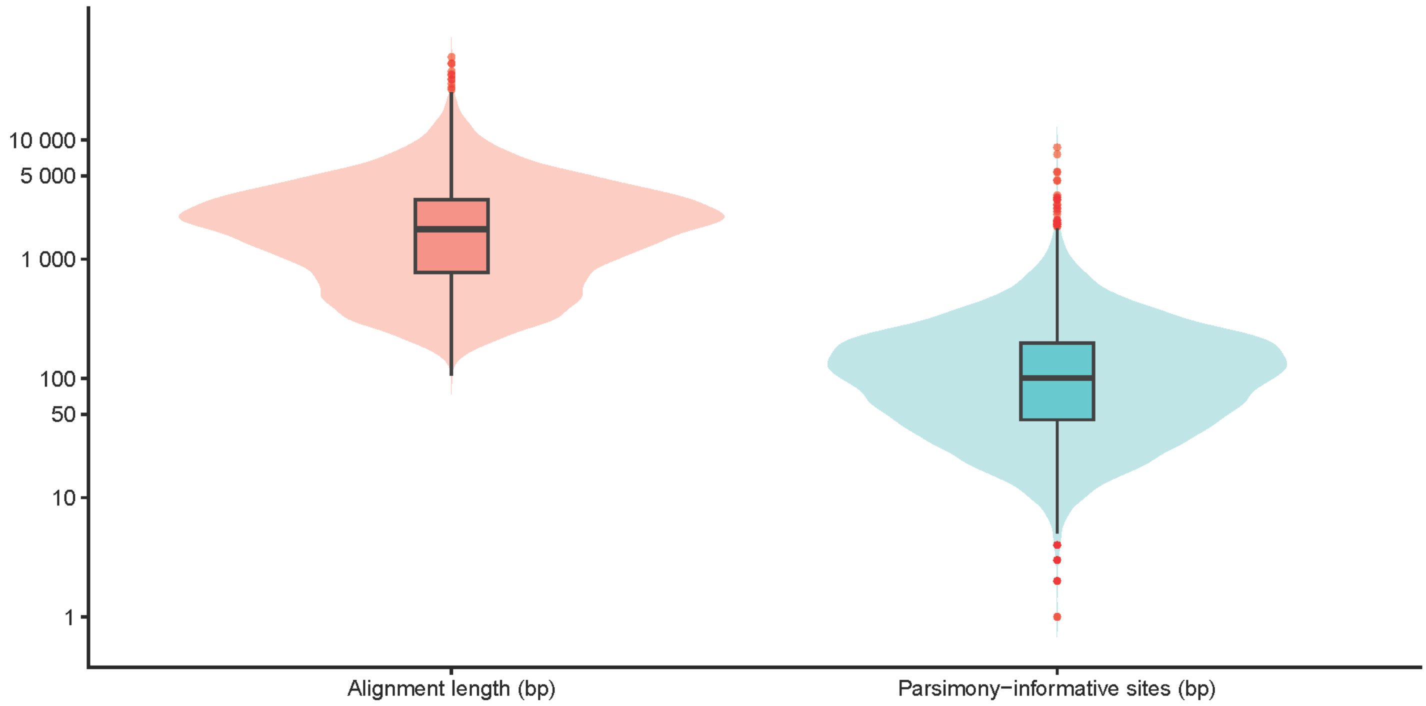


Fig. S1. Distribution of trimmed alignment length (bp) and parsimony-informative sites across 18,784 loci.

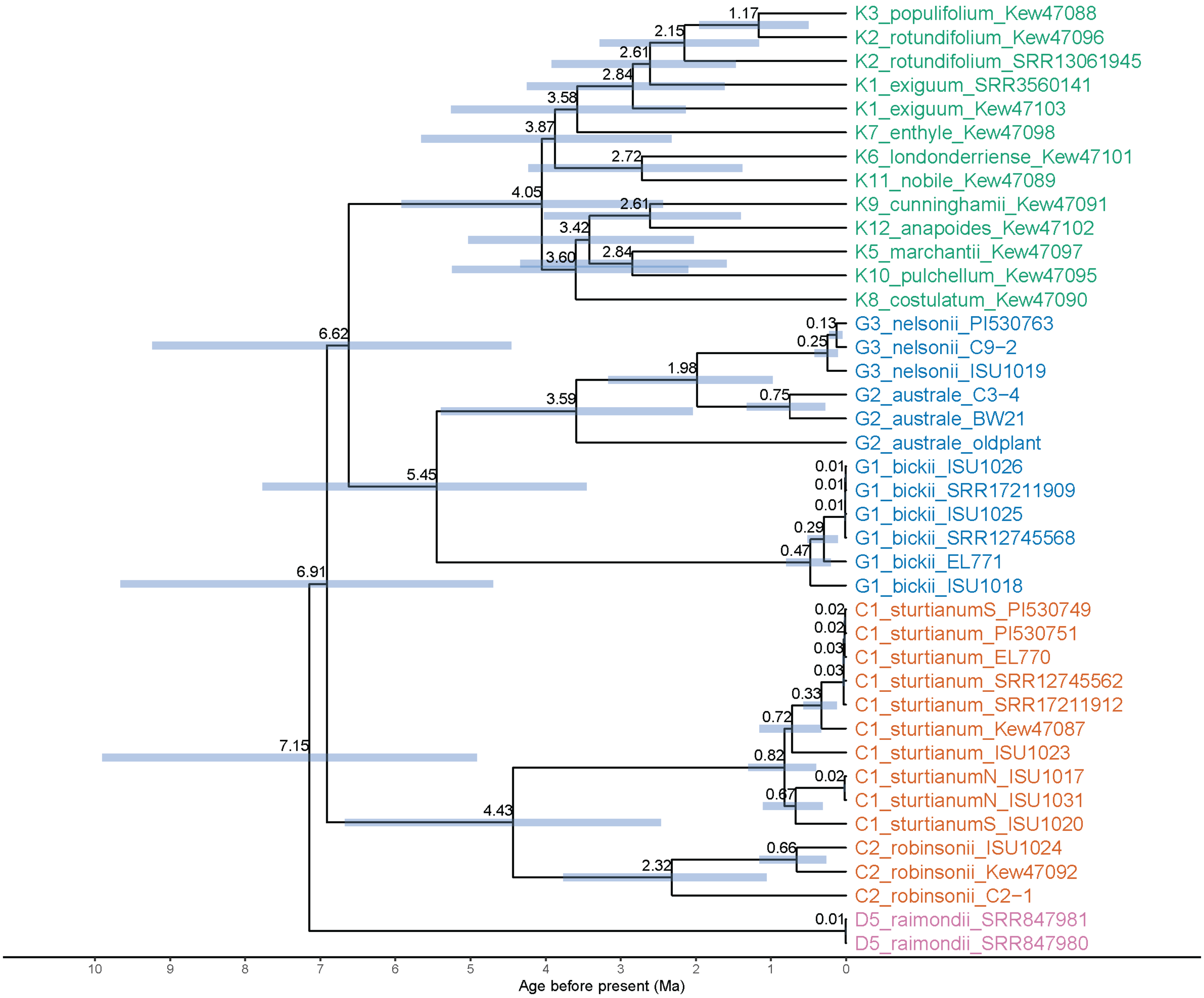


Fig. S2. Bayesian divergence-time estimates for Australian cottons. Node labels indicate divergence times between lineages, blue bars show the 95% corresponding intervals.

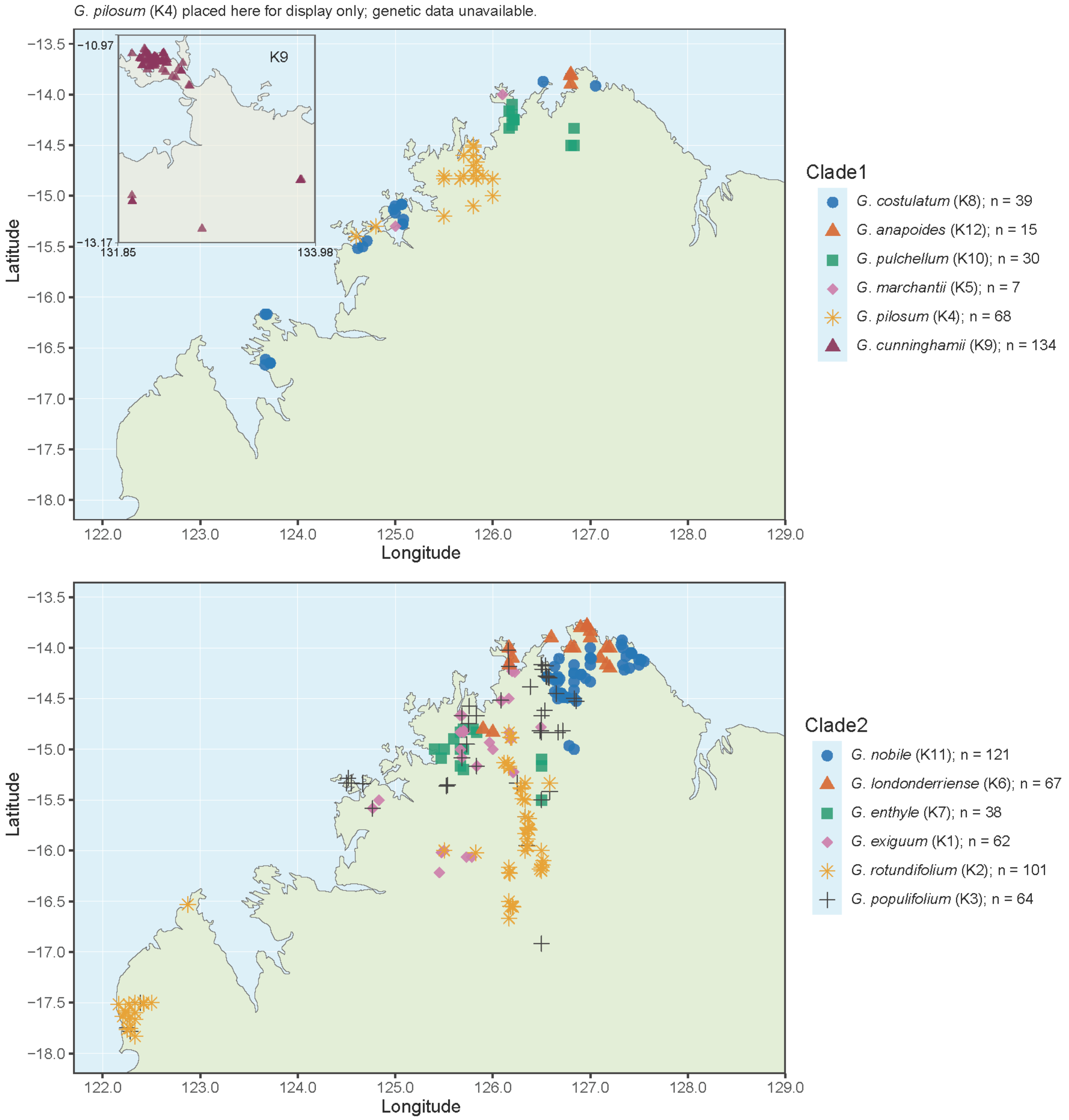


Fig. S3. Geographic distributions of the two clades of sect. *Grandicalyx* in Western Australia. Colors and symbols distinguish species.

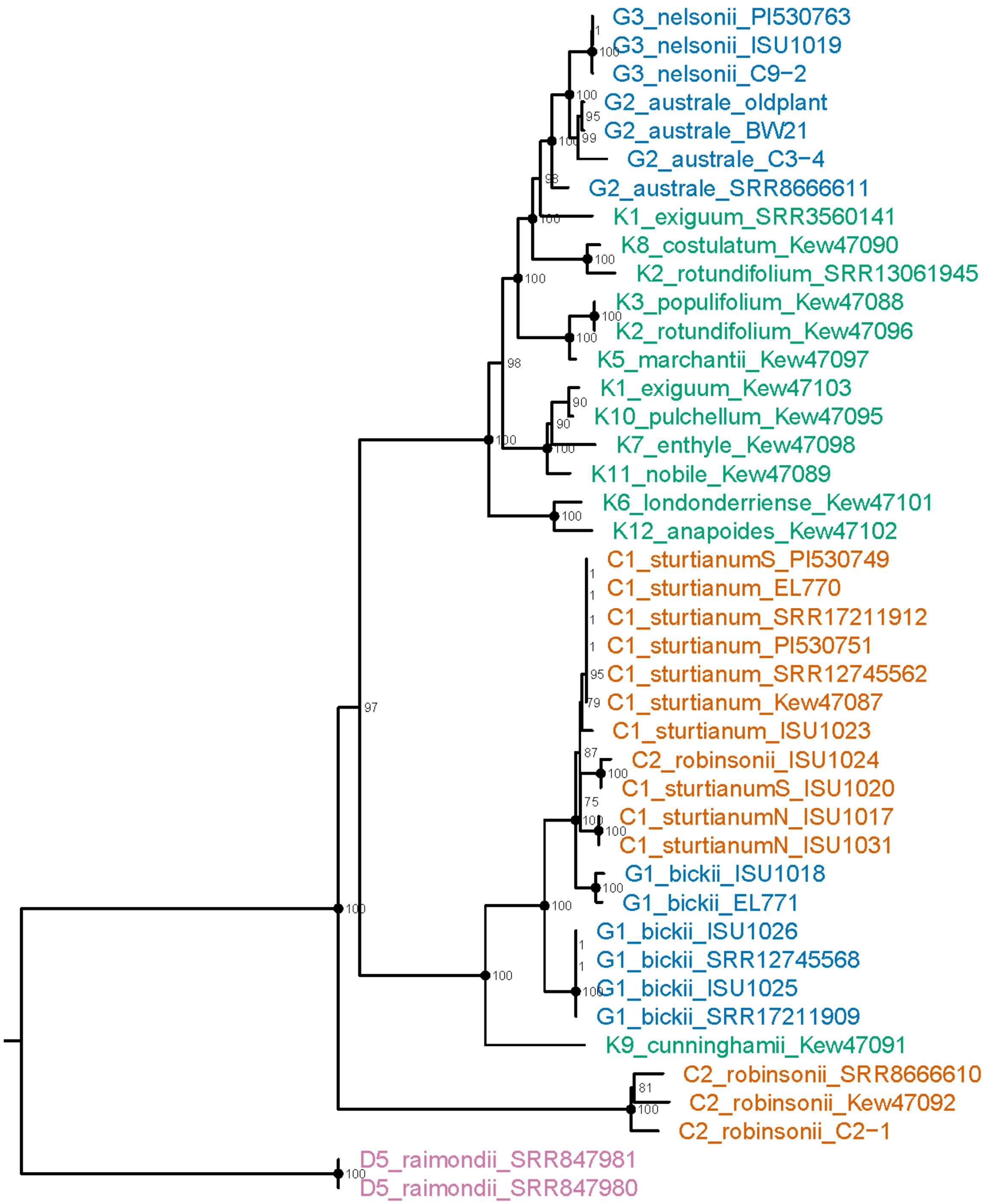


Fig. S4. Phylogeny inferred from chloroplast protein-coding genes, with bootstrap support values shown at internal nodes.

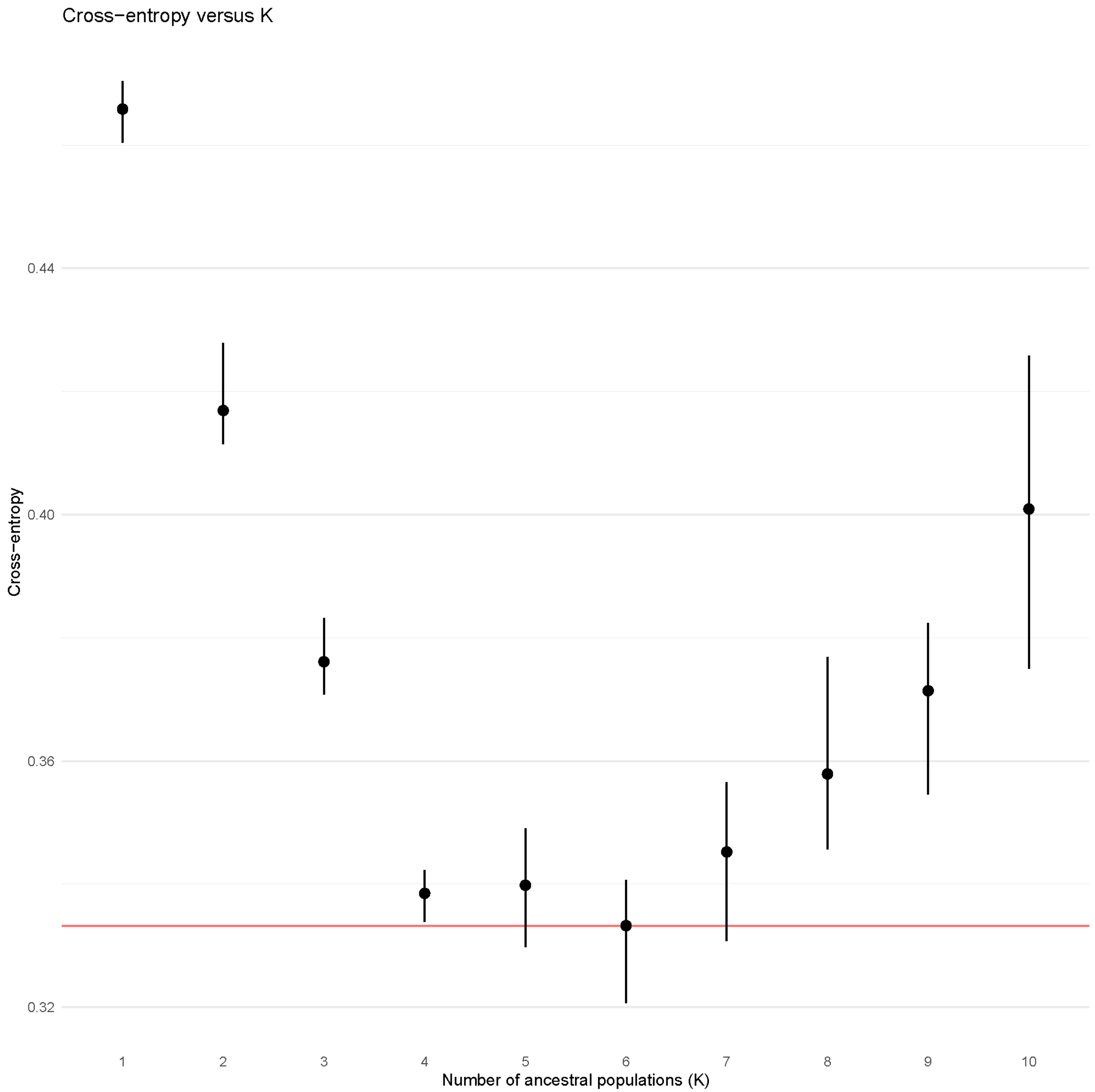


Fig S5. Cross-entropy values and the optimal number of ancestral populations.

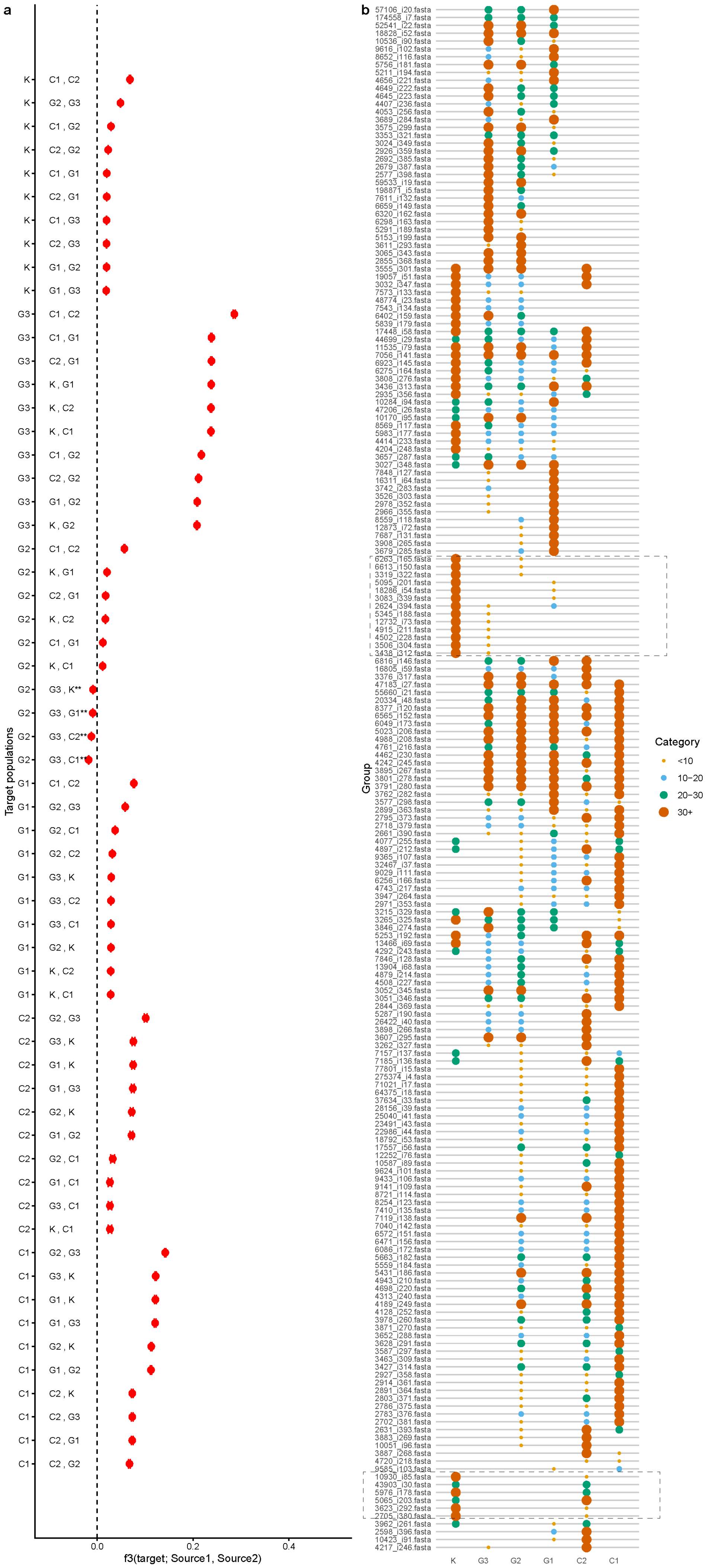


Fig S6. Genetic *f*3 admixture analysis (a) and KmerCity-based (b) genomic composition analysis of k-mer overlap patterns.
